# Safety and Efficacy of a Novel Glycoengineered Recombinant Vaccine Candidate against *Haemonchus contortus* in Sheep

**DOI:** 10.1101/2025.04.30.651318

**Authors:** Floriana Sajovitz-Grohmann, Isabella Adduci, Dirk Werling, Sandra Wiedermann, Licha N. Wortha, Bojan Prole, Julia Zlöbl, Jolina Elster, Alexander Tichy, Anja Joachim, Thomas Wittek, Barbara Hinney, Shi Yan, Katharina Lichtmannsperger

## Abstract

*Haemonchus contortus* is considered the most pathogenic nematode in small ruminants and South American camelids, causing significant production losses and threatening animal welfare worldwide. Compared to the use of anthelmintic drugs, to which resistance is increasing, vaccination is considered a more sustainable control strategy. The safety and efficacy of a novel glycoengineered vaccine produced in Hi5 insect cells was evaluated in a randomized, controlled vaccination-and-challenge trial. For this, 35 male Jura x Lacaune sheep were assigned to five groups (n=7). Three experimental groups were vaccinated three times subcutaneously with 1 ml of either the commercial Barbervax^®^ vaccine, the novel glycoengineered vaccine, consisting of a cocktail of five antigens (H11, H11-1, H11-2, H11-4 and GA1), or its non-glycoengineered counterpart before challenge with 5,000 *H. contortus* third-stage larvae. Clinical assessments, differential blood counts, serum antibody responses and fecal egg counts were evaluated at least once per week within the 16-week trial period. The abomasal worm burden was counted 40 days after challenge. The results of the mixed model showed a lower degree of anemia (mean packed cell volume (PCV) of 26.77%; mean hemoglobin (Hb) of 9.03 g/dl), in sheep vaccinated with the glycoengineered antigen cocktail compared to the unvaccinated group (PCV of 24.34%; mean Hb of 7.84 g/dl) and to the sheep vaccinated with the non-glycoengineered antigens (PCV of 26.11%; mean Hb of 8.53 g/dl). However, the degree of anemia was lower in sheep vaccinated with native antigens (Barbervax^®^) (PCV of 29.10%; mean Hb of 9.84 g/dl). Serum IgG and IgE antibodies were elevated in all vaccinated groups. Sheep vaccinated with the glycoengineered antigens showed a reduction in abomasal worm burden of 25.36% and a reduction of fecal egg shedding of 81.09%, the group vaccinated with non-glycoengineered antigen showed no worm reduction and a fecal egg count reduction of 32.72%. While there was no significant difference in egg shedding between sheep vaccinated with the glycoengineered antigens and sheep vaccinated with Barbervax^®^ (98.32%; p = 0.33), the Barbervax^®^ group showed a greater reduction in worm burden (86.40%). The results emphasize that glycoengineering of vaccine candidates is essential to achieve protective immunity against *H. contortus* in sheep and that the novel glycoengineered vaccine produced in insect cells clearly has the potential to reduce fecal egg shedding and worm burden, paving the way for the development of highly effective recombinant vaccines against this and other parasitic worms in the future.

## Introduction

*Haemonchus contortus* is the most pathogenic gastrointestinal nematode in small ruminants and South American camelids. Infections cause significant production losses (decreased weight gain and fertility and loss of young stock) and threaten animal welfare worldwide, especially in tropical and subtropical areas^1–6^. As a single worm ingests up to 50 μl of blood per day, even moderate infections can cause severe blood loss resulting in anemia and hypoproteinemia, and in severe cases even death may occur^5,7^. Currently, parasite control primarily relies on the administration of anthelmintic drugs, which has several disadvantages, including unwanted ecotoxicological effects, long withdrawal periods for meat and milk and an increasing prevalence of anthelminthic resistant worms^8–10^. Due to a lack of treatment response of resistant *H. contortus,* clinical disease will progress in affected hosts^10–13^.

In contrast, vaccination is considered the most promising strategy to control the parasite in susceptible domestic animal species. In the past a repertoire of potential vaccine targets, including somatic and secreted antigens, was identified and evaluated in *in vivo* trials^14^. Among the evaluated protein-based vaccine candidates, the H11 aminopeptidases and the H-gal-GP protease complex showed the highest efficacy^15,16^. Sheep vaccinated with either the native H11 or H-gal-GP antigens showed significant reductions in fecal egg shedding of 72%-93% and of 68%-69% in worm burden^17,18^. Both, H11 and H-gal-GP are “hidden” proteins, located in the intestine of adult worms, and are not naturally exposed to the host’s immune system during worm infection. Thus, a vaccine response to these antigens is not boosted by infection, and repeated immunization is needed to maintain a protective effect during the grazing season^19^. H11 and H-gal-GP are major components of the only commercial vaccine currently available against *H. contortus* – the Barbervax^®^ which was licensed in Australia in 2014 and is available in only a few countries.

To meet local biosecurity regulations, Barbervax^®^ is manufactured at the Albany Laboratory of the Department of Agriculture and Food in Western Australia from adult *Haemonchus* worms harvested on an industrial scale from the stomach of donor sheep after slaughter. One production step of Barbervax^®^ involves mechanical disruption of the worms, by which 480 mg of antigen can be obtained from 90 g of worms^20,21^. For commercial scale production, machines were implemented to open the stomachs and harvest the worms, antigens are then isolated and formulated as a vaccine^20^. This is currently the only way to obtain adult worms for antigen preparation, as *in vitro* parasite culture systems for the propagation and multiplication of *H. contortus* have so far not been established^22^. Attempts to produce recombinant H11 vaccine antigens, using conventional expression platforms or the free-living nematode *Caenorhabditis elegans,* have been reported previously^23–26^. The resulting products were unable to reach a satisfactory efficacy regarding worm burden and/or fecal egg count^14^. H11 proteins are naturally glycosylated, carrying very distinct N-glycan structures compared to those found in mammals. Such glycan modifications are often considered crucial for maintaining the antigenicity^25,27^. To address this, a novel glycoengineered recombinant vaccine was developed, comprising a cocktail of five soluble proteins (H11, H11-1, H11-2, H11-4 and GA1) produced using Hi5 insect cells, designed to incorporate nematode-type glycan modifications and enhance solubility^28^.

The objectives of the present study were to compare hematological and parasitological parameters in sheep following vaccination with the novel recombinant vaccines in comparison to a Barbervax^®^-vaccinated group (BVAX), an infected, unvaccinated (POS) and an uninfected, unvaccinated (NEG) control. It was hypothesized that vaccination of sheep with the novel vaccine and their subsequent challenge would result in reduced anemia, significant decrease in fecal egg shedding and abomasal worm burden and an elevated concentration of specific immunoglobulins in comparison to the positive control group, that results are comparable or better than those seen in sheep of the BVAX group and that the level of protection is higher in sheep vaccinated with the glycoengineered form (GEA group) of the vaccine compared to the non-glycoengineered form (NEA group).

## Results

### Clinical assessment

The clinical anemia scoring (FAMACHA^©^) of the BVAX, the GEA, the NEA and the negative control group (NEG) was significantly lower than in the positive control group (POS) after challenge (≥ week 10) (p<0.01). The mixed model analysis showed a mean FAMACHA^©^ score of 2.42 for NEG (95%CI = 2.26; 2.59), 3.46 for POS (95%CI = 3.29; 3.63), 2.27 for BVAX (95%CI = 2.10; 2.44), 2.31 for GEA (95%CI = 2.14; 2.48) and 2.17 for NEA group (95%CI = 2.00; 2.33) (**Fig. 1**).

**Figure 1:**
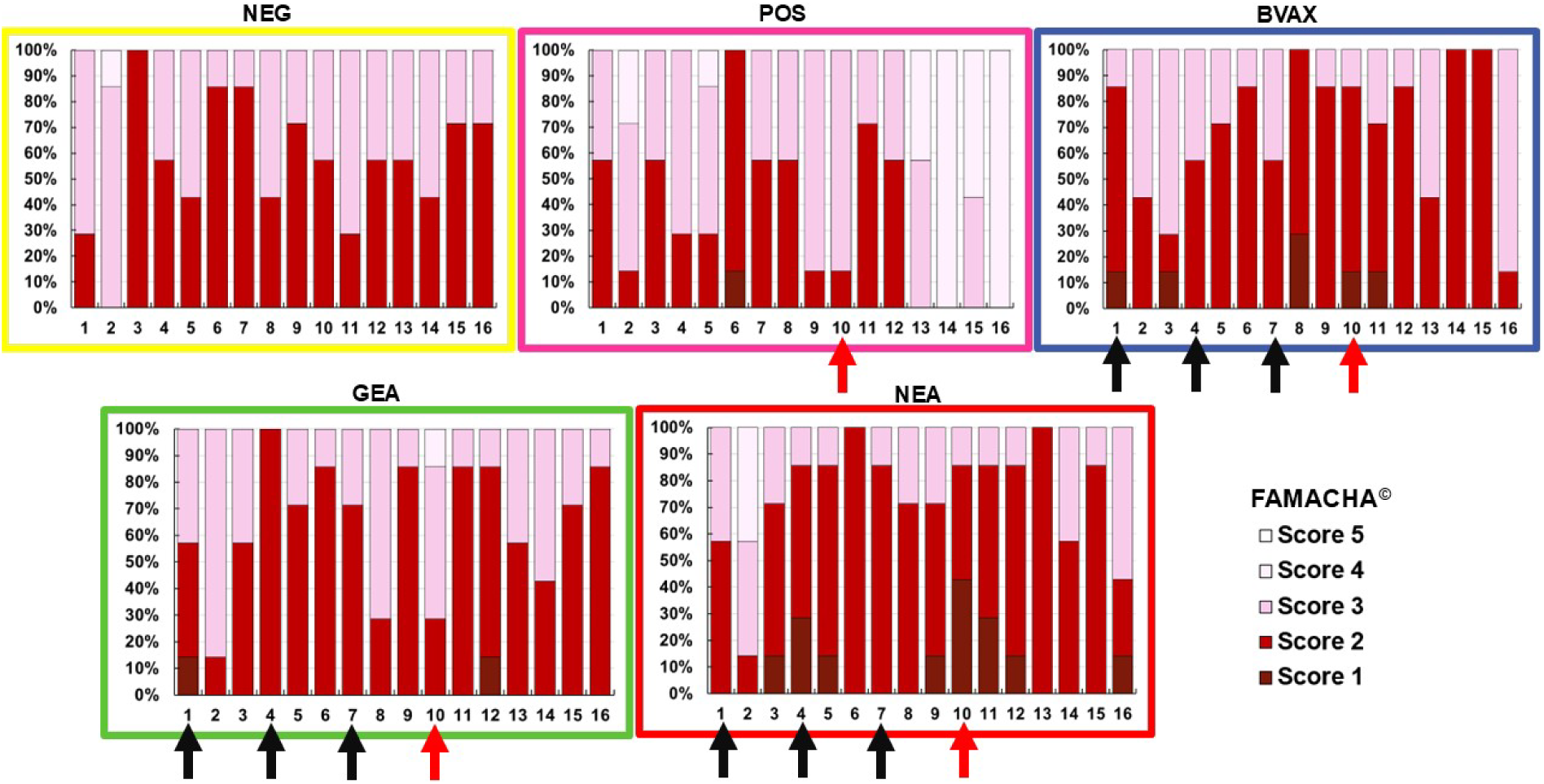
FAMACHA^©^ scores in the five study groups (n=7 sheep/group) during the 16-week study period. The five graphs show the FAMACHA^©^ results for the NEG (yellow; no vaccine, no challenge), the POS (pink; no vaccine but challenge with 5,000 *H. contortus* L3), the BVAX (blue; sheep vaccinated with Barbervax^®^ and challenged with 5,000 *H. contortus* L3), GEA (green; sheep vaccinated with the glycoengineered recombinant vaccine and challenged with 5,000 *H. contortus* L3) and NEA (red, sheep vaccinated with the non-glycoengineered recombinant vaccine and challenged with 5,000 *H. contortus* L3) groups. Each column shows the relative percentage of the results of the FAMACHA^©^ score of the seven sheep within the group in study week 1 to study week 16. The black arrows highlight the 1^st^ (week 1), 2^nd^ (week 4) and 3^rd^ (week 7) vaccination with either BVAX, GEA or NEA. The red arrows highlight week 10, when the sheep were challenged with 5,000 L3 *H. contortus*.

In addition to the FAMACHA^©^ score, changes in daily body weight gain (DBWG) were evaluated. Prior to L3 challenge (week 1 to 9; 63 days in total) no differences between the groups in DBWG, which ranged from 95 g to 222 g between the groups, were observed (**Fig. 2**; p = 0.87). After challenge (≥ week 10; 40 days in total), there was one significant difference between the BVAX and the NEG control group with sheep in BVAX group gaining less weight (**Fig. 2**; p = 0.02). All other groups showed a numerical but no statistical difference in DBWG. The NEG group gained a median of 213 g/d (min of 138 g, max of 238 g), the POS 150 g/d (min of 125 g, max of 188 g), the BVAX 138 g/d (min of 88 g, max of 138 g), the GEA 150 g/d (min of 113 g, max of 225 g) and the NEA 175 g/d (min of 100 g, max of 225 g).

**Figure 2:**
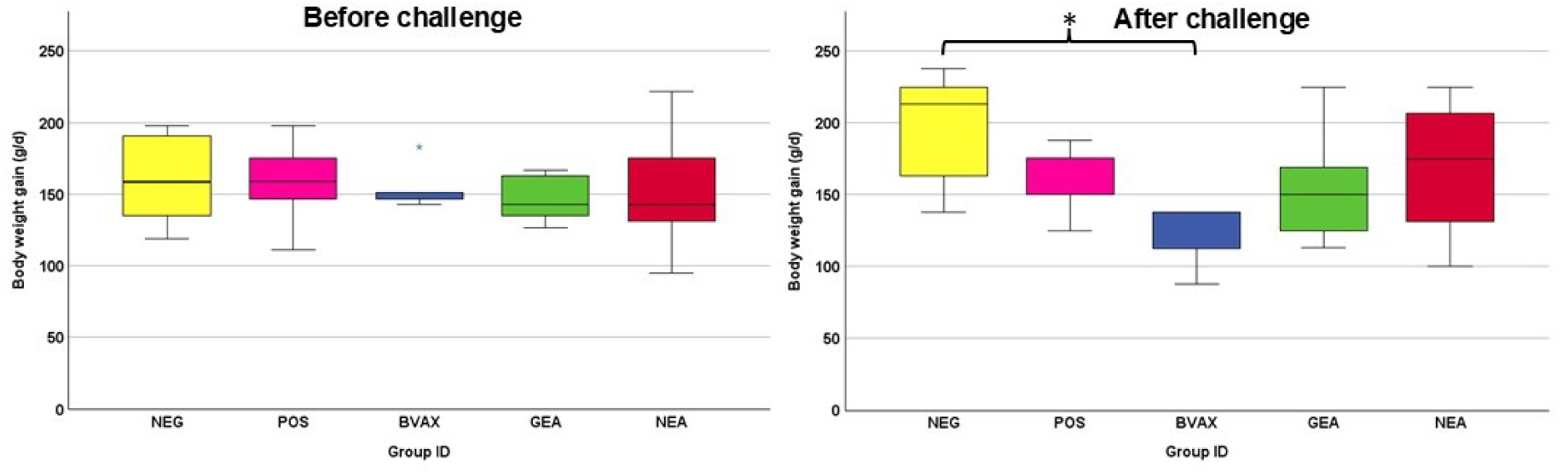
Overview on the daily body weight gain before and after challenge.

### Both GEA and Barbervax^®^ improve the values of hematological parameters

Hematological parameters, including packed cell volume (PCV), erythrocytes per milliliter and hemoglobin concentrations, were measured as indicators of potential blood loss and anemia. All three parameters were lowest in the POS group and highest in the NEG group after challenge (**Fig. 3** and **Table 1**).

**Figure 3:**
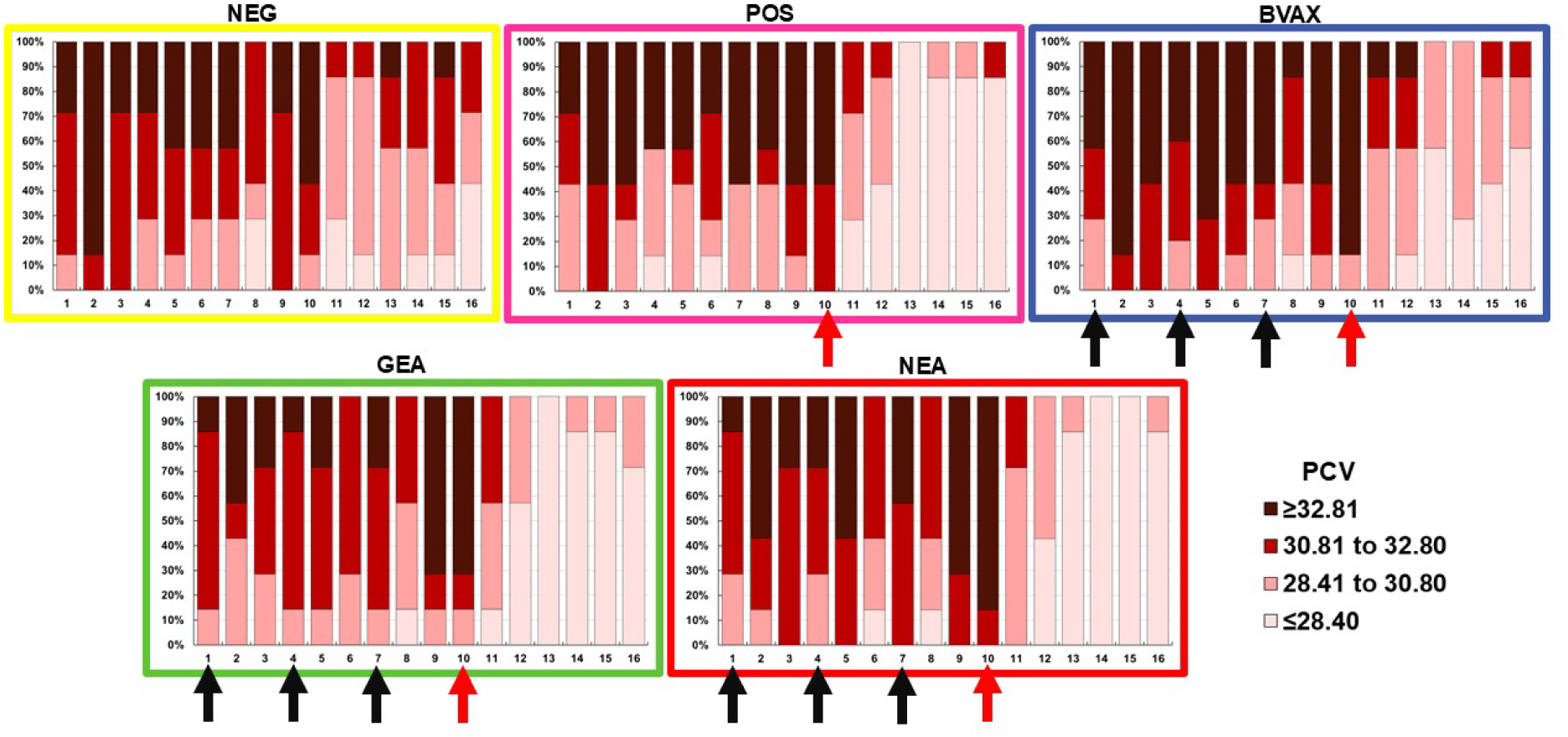
Packed cell volume (PCV [%]) in the five study groups over the study period of 16 weeks. For color code details see caption Fig. 1. The five graphs show the PCV results of the NEG, POS, BVAX, GEA and NEA group. Each column shows the relative percentage of the results of the PCV of the seven sheep within the group in study week 1 to study week 16. The black arrows highlight the 1^st^ (week 1), 2^nd^ (week 4) and 3^rd^ (week 7) vaccination with either BVAX, GEA or NEA. The red arrows highlight week 10, when the sheep were challenged with 5,000 L3 *H. contortus*.

**Table 1:**
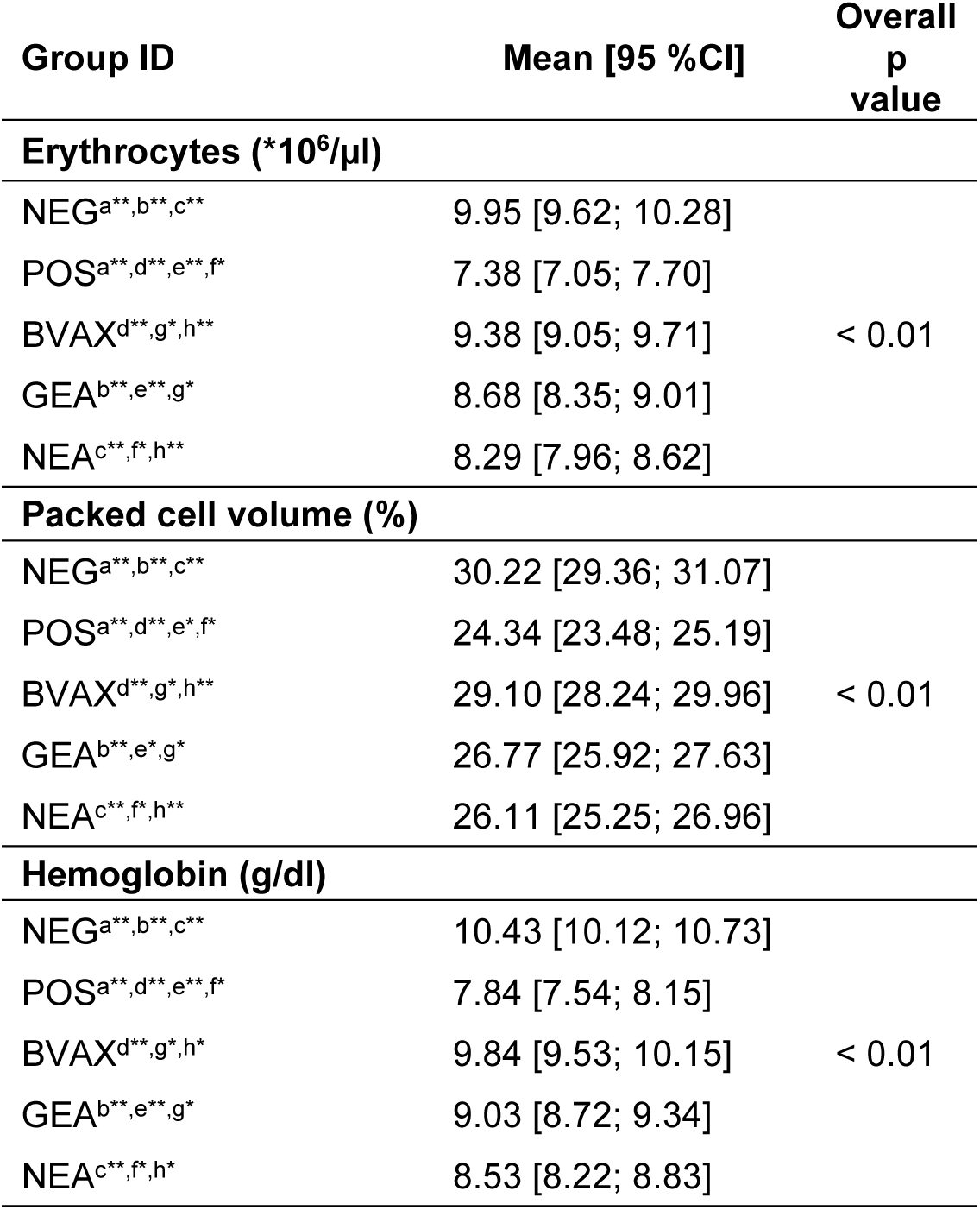
Differences between the five groups (indicated by superscript letters) in hematological parameters (number of erythrocytes per microliter, packed cell volume, hemoglobin concentration) after challenge (≥ week 10) assessed by Advia^®^ 2120i (Siemens, Erlangen, Germany). Superscript letters highlighted with * indicate p values ≤ 0.05, those with ** indicate p values < 0.01 for differences between groups with the same letter. The p values in the right column indicate differences between the groups overall.

| Group ID | Mean [95 %CI] | Overall<br>p<br>value |
| --- | --- | --- |
| Erythrocytes (*10<sup>6</sup>/μl) |  |  |
| NEG<sup>a**,b**,c**</sup> | 9.95 [9.62; 10.28] | < 0.01 |
| POS<sup>a**,d**,e**,f*</sup> | 7.38 [7.05; 7.70] |  |
| BVAX<sup>d**,g*,h**</sup> | 9.38 [9.05; 9.71] |  |
| GEA<sup>b**,e**,g*</sup> | 8.68 [8.35; 9.01] |  |
| NEA<sup>c**,f*,h**</sup> | 8.29 [7.96; 8.62] |  |
| Packed cell volume (%) |  |  |
| NEG<sup>a**,b**,c**</sup> | 30.22 [29.36; 31.07] | < 0.01 |
| POS<sup>a**,d**,e*,f*</sup> | 24.34 [23.48; 25.19] |  |
| BVAX<sup>d**,g*,h**</sup> | 29.10 [28.24; 29.96] |  |
| GEA<sup>b**,e*,g*</sup> | 26.77 [25.92; 27.63] |  |
| NEA<sup>c**,f*,h**</sup> | 26.11 [25.25; 26.96] |  |
| Hemoglobin (g/dl) |  |  |
| NEG<sup>a**,b**,c**</sup> | 10.43 [10.12; 10.73] | < 0.01 |
| POS<sup>a**,d**,e**,f*</sup> | 7.84 [7.54; 8.15] |  |
| BVAX<sup>d**,g*,h*</sup> | 9.84 [9.53; 10.15] |  |
| GEA<sup>b**,e**,g*</sup> | 9.03 [8.72; 9.34] |  |
| NEA<sup>c**,f*,h*</sup> | 8.53 [8.22; 8.83] |  |

Globulin and total protein concentrations were higher in the GEA group in comparison to the POS group depending on the time point after challenge (**Fig. 4**). There were higher globulin concentrations (calculated as total protein minus albumin) in the BVAX and the GEA group in comparison to the NEA group over the whole period. There were no differences in total protein between all groups in week 8 (p = 0.08), week 9 (p = 0.06) and week 10 (p = 0.06). One week post challenge (week 11), POS was significantly lower in total protein than the BVAX (p < 0.01), GEA (p < 0.01) and NEA (p = 0.02). In week 12, there were no differences between the groups. In week 13, there was a difference between the POS and NEG group (p < 0.01) and between BVAX and the POS (p < 0.01). The NEA group showed significantly lower total protein concentrations than the NEG group. In week 14, BVAX and GEA showed significantly higher total protein concentrations than the POS group (p = 0.01). In week 15, BVAX and GEA groups showed significantly higher total protein concentrations than the POS (p = 0.01) and the NEG group showed higher total protein concentrations than the POS (p = 0.01). In week 15, there was a significant difference between the POS and NEG group (p = 0.04) and between the BVAX and the POS group (p = 0.05).

**Figure 4:**
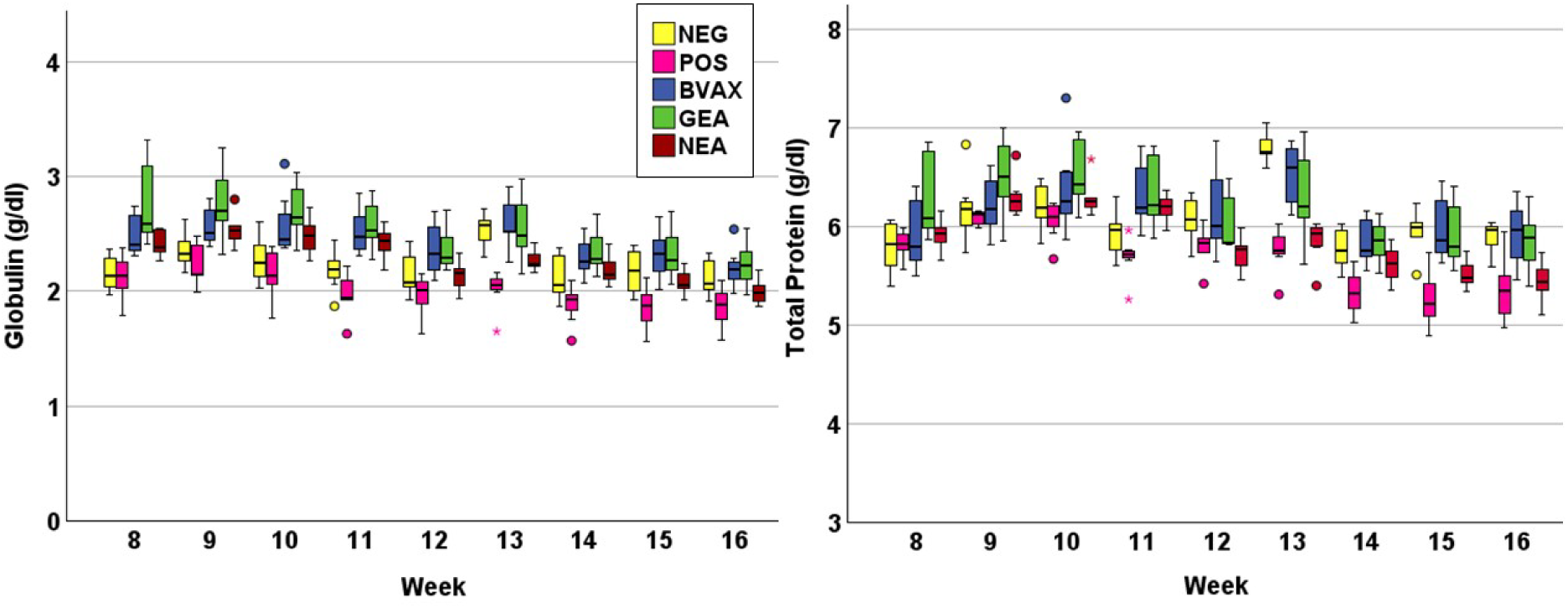
Globulin and total protein concentrations in serum in g/dl before (< week 10) and after (≥ week 10) challenge. Sheep in the POS (pink), BVAX (blue), GEA (green) and NEA (red) group were challenged with 5,000 *H. contortus* L3 in week 10.

The results were underscored by statistical evaluation including the timepoints ≥ week 10 (after challenge) where there was a clear difference in total protein between the GEA and the POS group (GEA: mean of 5.95 g/dl; POS mean of 5.58 g/dl) and between GEA and NEA (GEA: mean of 5.95 g/dl; NEA: mean of 5.58 g/dl) (**Tab. 2**). There was no difference between BVAX and GEA in albumin concentrations, but between BVAX (mean of 3.76 g/dl) and NEA (mean of 3.52 g/dl) a difference was seen (p ≤ 0.05) (**Tab. 2**). In week 10 (p = 0.48) and week 11 (p = 0.79) there were no differences in albumin concentrations. In week 12, there was one difference between the NEA and the NEG group (p = 0.02). In week 13, there was a difference between the NEA (p < 0.01) and GEA (p = 0.05) and the NEG, the POS and the NEG group (p = 0.01). In week 14, there were no differences. In weeks 15 and 16, there were differences between the POS and the NEG control group (p = 0.01 resp. p = 0.05) and the NEA and the NEG group (p = 0.02 resp. p = 0.02).

**Table 2:**
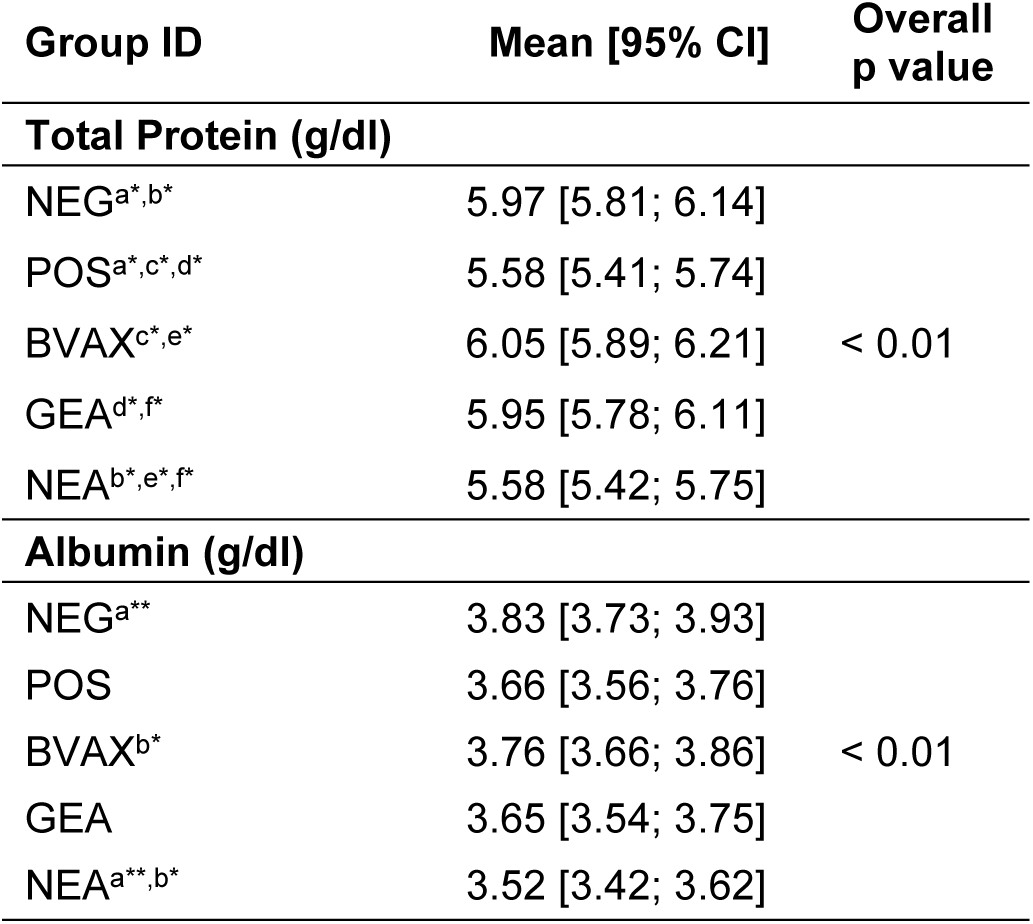
Differences in total protein and albumin between the five groups (7 sheep per group) after challenge in week 10. All superscript letters highlighted by * indicate p values ≤ 0.05. Superscript letters with ** indicate p values < 0.01. The p values in the right column indicate differences between the groups overall.

Differences in segmented neutrophils, eosinophils, basophile granulocytes and lymphocytes were detected between the groups. Statistically significant differences were seen in the number of eosinophils, with the BVAX group showing the highest numbers throughout the trial. The relative distribution of the white blood cells per group over the period of 16 weeks is shown in **Fig. 5**. Notably, all vaccinated groups showed a marked increase of eosinophils after the second vaccination (week 5). Two weeks after the challenge infection (week 12), increased eosinophils appeared in the POS control group, BVAX and GEA groups but not the NEA group. The statistical evaluation showed that the sheep in GEA showed the highest number of lymphocytes (4679.40/µl), the sheep in the BVAX the highest number of eosinophils (507.17/µl) and basophils (52.75/µl) (**Tab. 3**).

**Figure 5:**
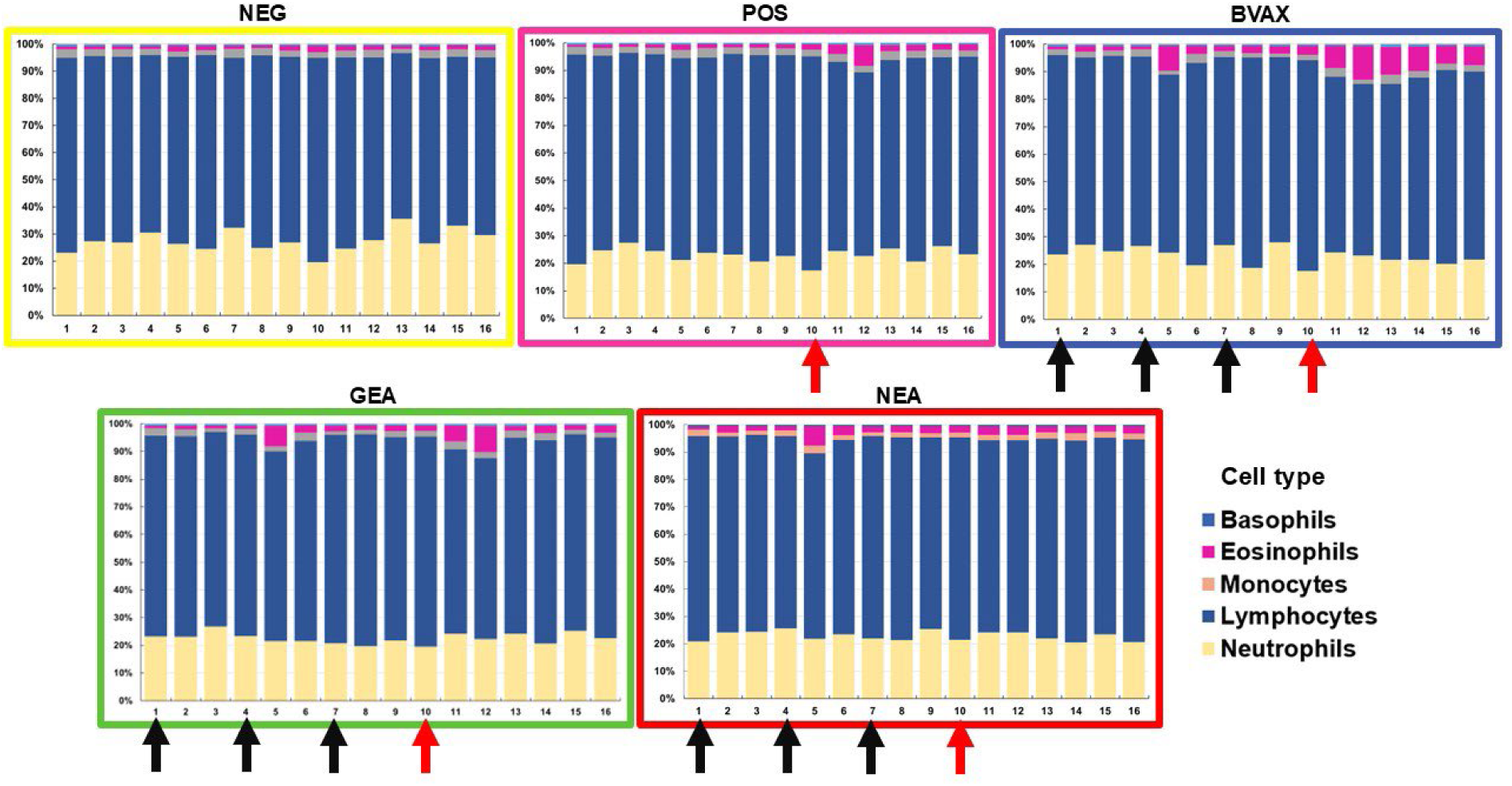
Relative distribution of granulocytes (segmented neutrophils, eosinophils, basophils), monocytes and lymphocytes of the five groups (for color code details see caption Fig. 1). In week 1 (p = 0.37), week 2 (p = 0.07), week 3 (p = 0.24) and week 4 (p = 0.54) there were no differences in eosinophil percentage between the five groups. Following, in week 5 (p = 0.00), week 6 (p = 0.01), week 7 (p = 0.04), week 8 (p = 0.01) there were differences between the groups. In week 9 (p = 0.08) and week 10 (p = 0.43) there were no differences in eosinophil percentages. From week 10 onwards there were differences in the eosinophil percentages between the groups (p ≤ 0.05). The black arrows highlight the 1^st^ (week 1), 2^nd^ (week 4) and 3^rd^ (week 7) vaccination with either BVAX, GEA or NEA. The red arrows highlight the week when the sheep were challenged with 5,000 L3 *H. contortus*.

**Table 3:** Differences in the differential blood counts between the five groups. Mean values and 95% confidence intervals are shown, the significance level was set at p ≤ 0.05. All superscript letters highlighted by * indicate p values ≤ 0.05. Superscript letters with ** indicate p values < 0.01. The p values in the right column indicate differences between the groups overall.

| Group ID | Mean [95 %CI] | Overall<br>p value |
| --- | --- | --- |
| Lymphocytes /μl |  |  |
| NEG<sup>a*</sup> | 4144.92 [3892.84; 4396.99] | < 0.01 |
| POS<sup>b**</sup> | 3890.61 [3638.54; 4142.68] |  |
| BVAX | 4391.28 [4139.21; 4643.35] |  |
| GEA<sup>a*,b**,c*</sup> | 4679.40 [4427.33; 4931.48] |  |
| NEA<sup>c*</sup> | 4004.36 [3752.28; 4256.43] |  |
| Monocytes /μl |  |  |
| NEG | 167.00 [139.39; 194.61] | 0.08 |
| POS | 143.44 [115.83; 171.05] |  |
| BVAX | 165.81 [138.21; 193.42] |  |
| GEA | 148.53 [120.92; 176.14] |  |
| NEA | 116.57 [88.96; 144.18] |  |
| Eosinophil Granulocytes /μl |  |  |
| NEG<sup>a**</sup> | 97.38 [28.44; 166.33] | < 0.01 |
| POS<sup>b**</sup> | 138.45 [69.50; 207.40] |  |
| BVAX<sup>a**,b**,c**,d**</sup> | 507.17 [438.22; 576.11] |  |
| GEA<sup>c**</sup> | 188.82 [119.87; 257.76] |  |
| NEA<sup>d**</sup> | 143.78 [74.83; 212.72] |  |
| Basophil Granulocytes /μl |  |  |
| NEG<sup>a**</sup> | 36.33 [31.22; 41.43] | < 0.01 |
| POS<sup>b**</sup> | 36.96 [31.85; 42.06] |  |
| BVAX<sup>a**,b**,c**</sup> | 52.75 [47.65; 57.86] |  |
| GEA | 42.82 [37.71; 47.93] |  |
| NEA<sup>c**</sup> | 33.78 [28.67; 38.88] |  |
| Segmented Granulocytes /μl |  |  |
| NEG<sup>a**,b**,c**,d**</sup> | 1874.23 [1755.93; 1992.52] | < 0.01 |
| POS<sup>a**</sup> | 1292.75 [1174.46; 1411.05] |  |
| BVAX<sup>b**</sup> | 1421.45 [1303.16; 1539.75] |  |
| GEA<sup>c**,e*</sup> | 1514.37 [1396.07; 1632.66] |  |
| NEA<sup>d**,e*</sup> | 1233.35 [1115.05; 1351.64] |  |

### All vaccines show similar impact on clinical parameters after the 1^st^, 2^nd^ and 3^rd^ vaccination

Body temperature was measured before vaccination and with a mean of 39.37 °C ± 0.23 °C there was no difference between the groups (p = 0.24). All sheep showed a bright and alert appearance and good appetite after vaccinations. After the second vaccination differences were seen on the day of vaccination (**Tab. 4**). At the day of the second vaccination one sheep vaccinated with GEA showed a body temperature of 39.6 °C 6 hours after vaccination and 41.5 °C 8.5 hours post vaccination and a slightly reduced demeanor.

**Table 4:** Mean body temperature and standard deviations in degree Celsius (°C) after vaccination with Barbervax^®^ (BVAX), GEA (glycoengineered recombinant vaccine) and NEA (non-glycoengineered recombinant vaccine). Body temperature was measured on the day of vaccination (day 0) and one day after the vaccination. P values show the results of the one-way ANOVA. * Highlights significant values.

| Vaccination | BVAX | GEA | NEA | p value |
| --- | --- | --- | --- | --- |
| 1 (day 0) | 39.86 ± 0.19 | 40.04 ± 0.46 | 39.93 ± 0.30 | 0.59 |
| 1 (day 1) | 39.64 ± 0.32 | 39.86 ± 0.35 | 39.76 ± 0.20 | 0.42 |
| 2 (day 0) | 39.54 ± 0.73 | 39.87 ± 0.57 | 40.36 ± 0.60 | 0.08 |
| 2 (day 1) | 39.33 ± 0.27 | 40.20 ± 0.66 | 39.80 ± 0.24 | 0.01* |
| 3 (day 0) | 39.03 ± 0.32 | 39.00 ± 0.17 | 39.10 ± 0.26 | 0.76 |
| 3 (day 1) | 39.45 ± 0.49 | 39.09 ± 0.52 | 39.33 ± 0.27 | 0.30 |

Induration, lymph node enlargement, swelling and pain was recorded in all experimental groups with differing degrees after each vaccination (for details see Supplementary File, Tables 1-4).

### The glycoengineered vaccine reduces egg shedding and eggs per female worm

To evaluate vaccine efficacy against infections with *H. contortus* fecal egg counts (displayed as EpG, eggs per gram of feces) and worm burden were assessed. The reduction of egg shedding was 98.32% for BVAX, 81.09% for GEA and 32.72% for NEA in comparison to the POS group (**Fig. 6**). BVAX showed lower egg counts than the POS group (p < 0.01) and the NEA group (p = 0.02). There was no difference between the BVAX and the GEA group (p = 0.33). GEA showed a numerical but not statistically significant lower egg shedding than the POS group (p = 0.08) and the NEA group (p = 1.0). The median cumulative EpG were 36,930 in the POS group, 620 in the BVAX group, 6,985 in the GEA group and 24,845 in the NEA group. The data of the egg shedding of the individual sheep is shown in **Fig. 6**.

**Figure 6:**
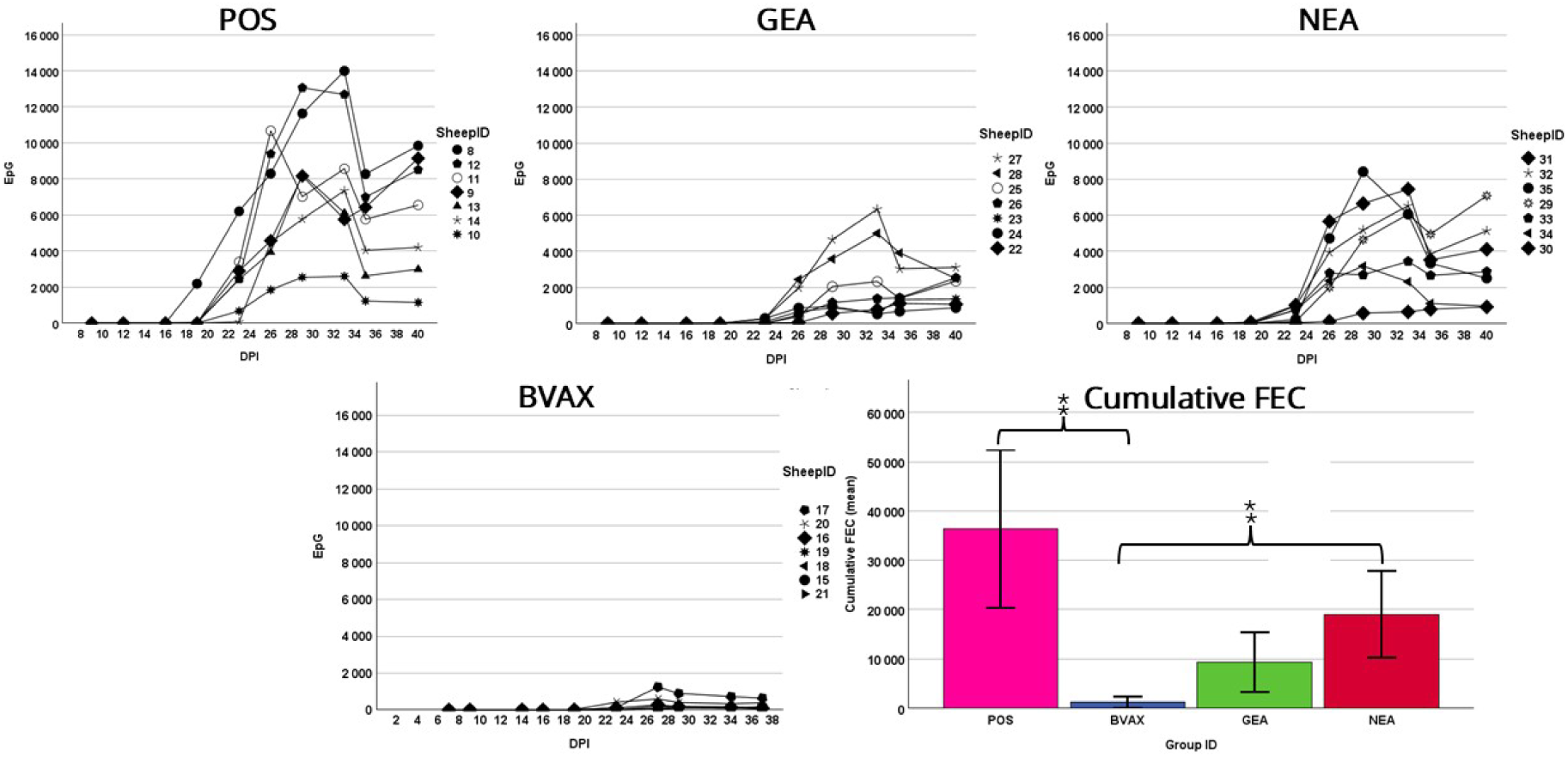
Individual fecal egg shedding (as eggs per gram of feces; EpG) after challenge with 5,000 *H. contortus* L3 (DPI: days post infection) of the POS group (not vaccinated but challenged sheep), the BVAX group, the GEA and the NEA group. The vaccinated sheep showed a delayed onset of egg shedding in comparison to the control group. Egg shedding in the POS group ranged from 10,005 to 60,425 EpG, in the BVAX 340 to 3,630 EpG, in the GEA 3,580 to 19,365 EpG and in the NEA 3,080 to 28,480 EpG. The error bars indicate the 95% CI.

In comparison to the POS group, the highest worm burden reduction was observed in the BVAX group (86.40%), whereas the GEA group had a lower reduction (25.40%), and the NEA group showed no reduction. There were significant differences between the BVAX and the GEA group (p = 0.04), between the BVAX and the POS (p < 0.01) and BVAX and NEA (p = 0.01). While the impact on the worm burden was not that pronounced, the number of eggs per female worm were significantly lower in GEA and BVAX groups compared to those of the POS group. There were differences between the POS and GEA (p < 0.01) and between the POS and the BVAX group (p < 0.01). GEA showed the overall lowest median value (7.98) (**Fig. 7)**.

**Figure 7:**
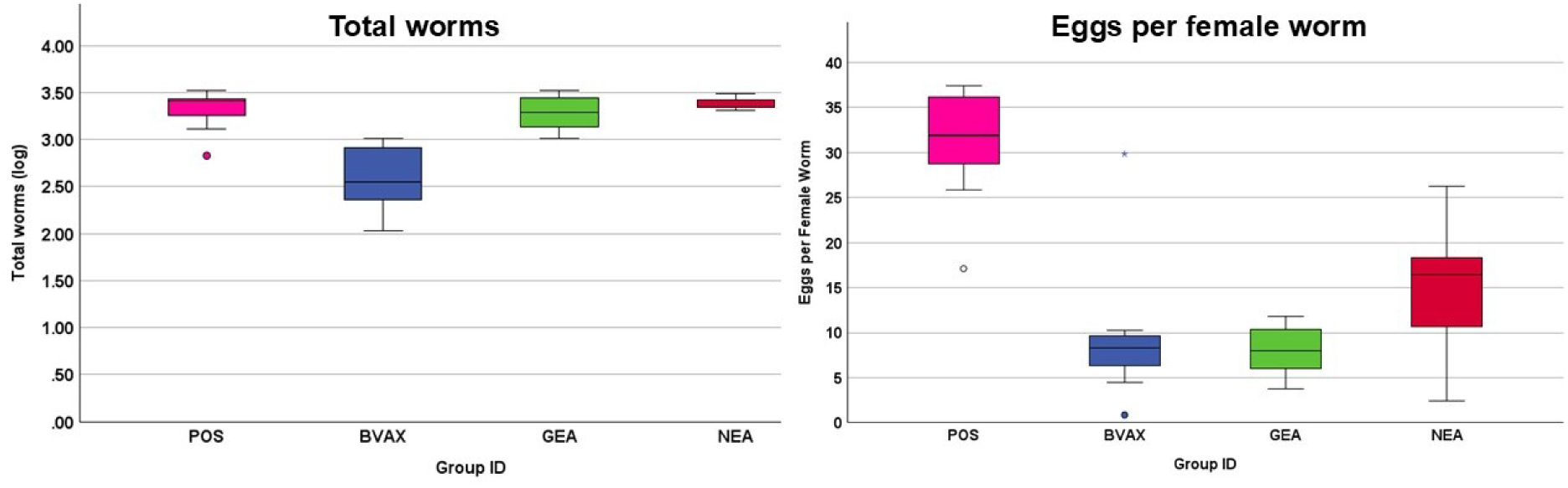
Number of eggs per female worm and worm burden. **Left**) Overview of the abomasal worm burden of 28 sheep euthanized 40 days after challenged with 5,000 *H. contortus* L3. **Right**) Ratio of eggs per female in the different groups.

In addition to these parameters, a possible impact of the vaccine on the establishment rate of ingested larvae was investigated. While this was 41.90% in the GEA and 48.44% in the NEA group and was lower (for GEA) that the 44.86% seen in the POS, the values did not reach the result seen for the BVAX group (10.28%).

The female: male ratio was 1.29 in the POS (min of 0.78, max of 1.67), 0.59 in the BVAX (min of 0.18, max of 1.02), 1.25 in the GEA (min of 0.82, max of 2.10) and 1.24 in the NEA group (min of 0.94, max of 1.48). The number of eggs per female worm were significantly lower in GEA and BVAX groups compared to those of the POS group (p < 0.01 for both comparisons)

### Animals in the GEA-vaccinated group show a faster increase of parasite- and vaccine-specific serum immunoglobulins

To determine whether vaccination with the recombinant antigens GEA and NEA elicited specific antibodies in animals, sheep sera collected during the trial were examined by ELISA. ELISA data initially indicated that the total serum IgG and IgM antibodies did not change among the groups and during the trial (**Fig. 8**). Subsequently, serum samples were analyzed using ELISA plates coated with either soluble proteins of adult *H. contortus* or with vaccine antigens (indirect ELISA). *Haemonchus*-specific IgG antibodies were induced in the vaccine groups (BVAX, GEA, NEA; **Fig. 9 A**). Vaccination with GEA and NEA led to a faster increase of IgG levels compared to sheep in the BVAX group which showed a more gradual antibody increase. Animals in the BVAX group maintained the highest IgG levels of all vaccine groups after the third vaccination, in line with the assumption that BVAX contains more than the described antigens. As seen for other parameters, vaccination of animals with GEA led to a higher level of IgG antibodies compared to antibodies induced by NEA. This pattern was also observed for *Haemonchus*-specific IgE antibodies (**Fig. 9 B**). No changes were detected for IgM antibodies in any group (**Supplementary Fig. 3**).

**Figure 8:**
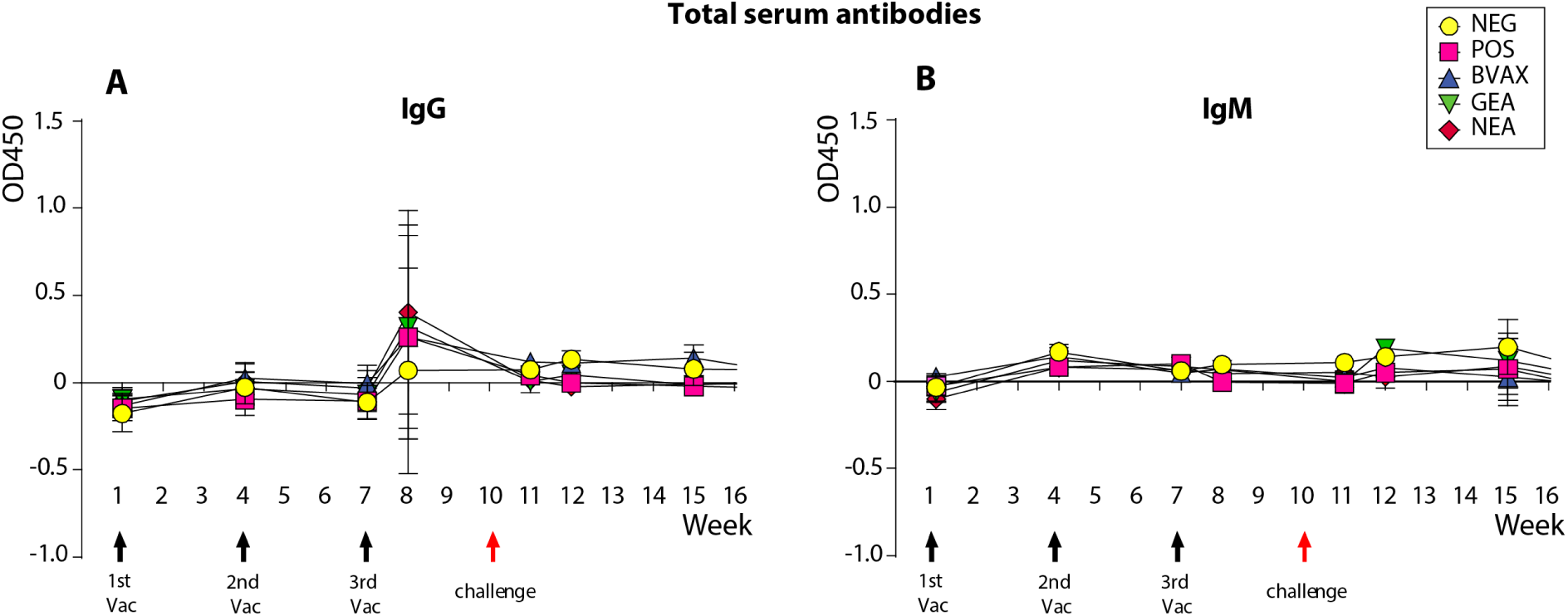
ELISA analysis of total serum antibodies in sheep. Panels show responses of baseline corrected-value total IgG (A) and IgM (B) antibodies, respectively. Assays were performed using plates coated with sheep sera.

**Figure 9:**
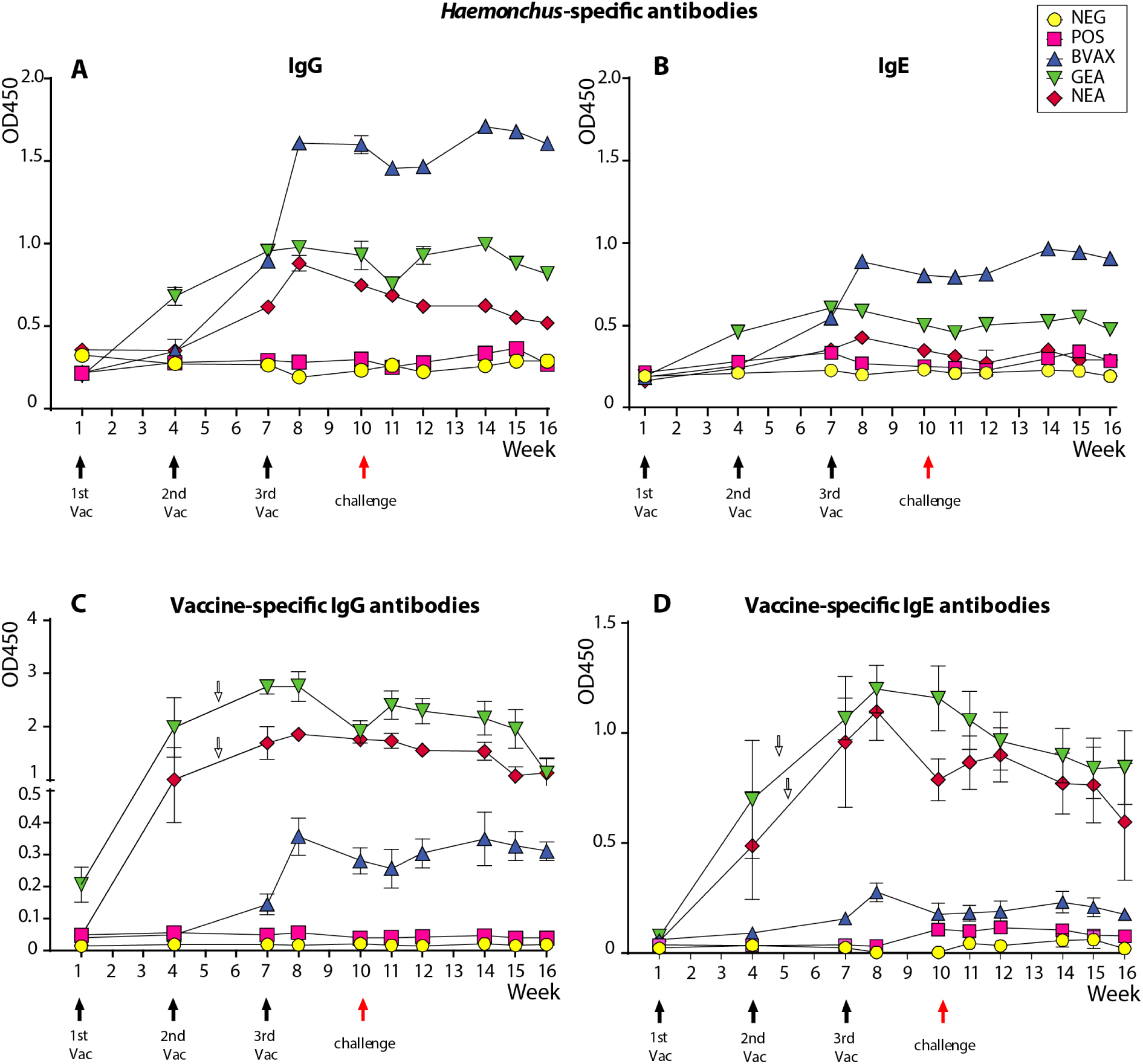
ELISA analysis of specific serum antibodies in sheep. Panels (A, B) show responses of *Haemonchus*-specific IgG and IgE antibodies, respectively. Assays were performed using plates coated with soluble proteins of adult parasites and combined sheep sera. Panels (C, D) illustrate antigen-specific IgG and IgE responses in individual sheep. Vaccine-specific antibodies for the GEA and NEA groups were measured using plates coated with an equal amount of the corresponding vaccine antigens (marked with white arrows); the BVAX and negative and positive control groups were assayed using Barbervax^®^-coated plates. Data are plotted on different y-axis scales to better visualize the pattern of the different assays.

To assess the immune responses of individual animals, vaccine-specific antibodies were assessed using plates coated with either the corresponding vaccine antigens, whereas for the control groups (NEG and POS) Barbervax^®^-coated plates were used. The results indicated elevated levels of antigen-specific IgG (**Fig. 9 C**) and IgE (**Fig. 9 D**) antibodies in all three vaccine groups (BVAX, GEA, NEA). Although less pronounced in the BVAX group compared to GEA and NEA groups, IgG and IgE specific to the respective vaccine antigens were induced post vaccination but slowly tapering off during the trial. Analysis of IgA and IgM antibodies did not reveal differences between the vaccine groups and control group; vaccine-specific salivary IgA antibodies were also measured using antigen-coated plates. However, no obvious changes

### Sera of GEA-vaccinated animals show a similar antigen-detection spectrum compared to Barbervax^®^-vaccinated animals

In a further set of experiments, the antigen recognition patterns of GEA-induced IgGs were compared to those seen in sera of Barbervax^®^-induced IgG by Western blotting. The first experiment demonstrated that IgGs in sera of the BVAX and GEA groups recognized a number of protein targets with a broad molecular weight range (20 – 250 kDa). These proteins were not only present in adult-stage parasite but also in the infective L3 larvae (**Fig. 10A**). This suggested a broader antibody targeting upon infection. *Haemonchus*-specific IgG antibodies in the sera of the GEA vaccinated group were detectable after the 1^st^ vaccination (week 4), presumably a few weeks earlier than the appearance of IgGs in sera of sheep vaccinated with Barbervax^®^ (BVAX group).

**Figure 10:**
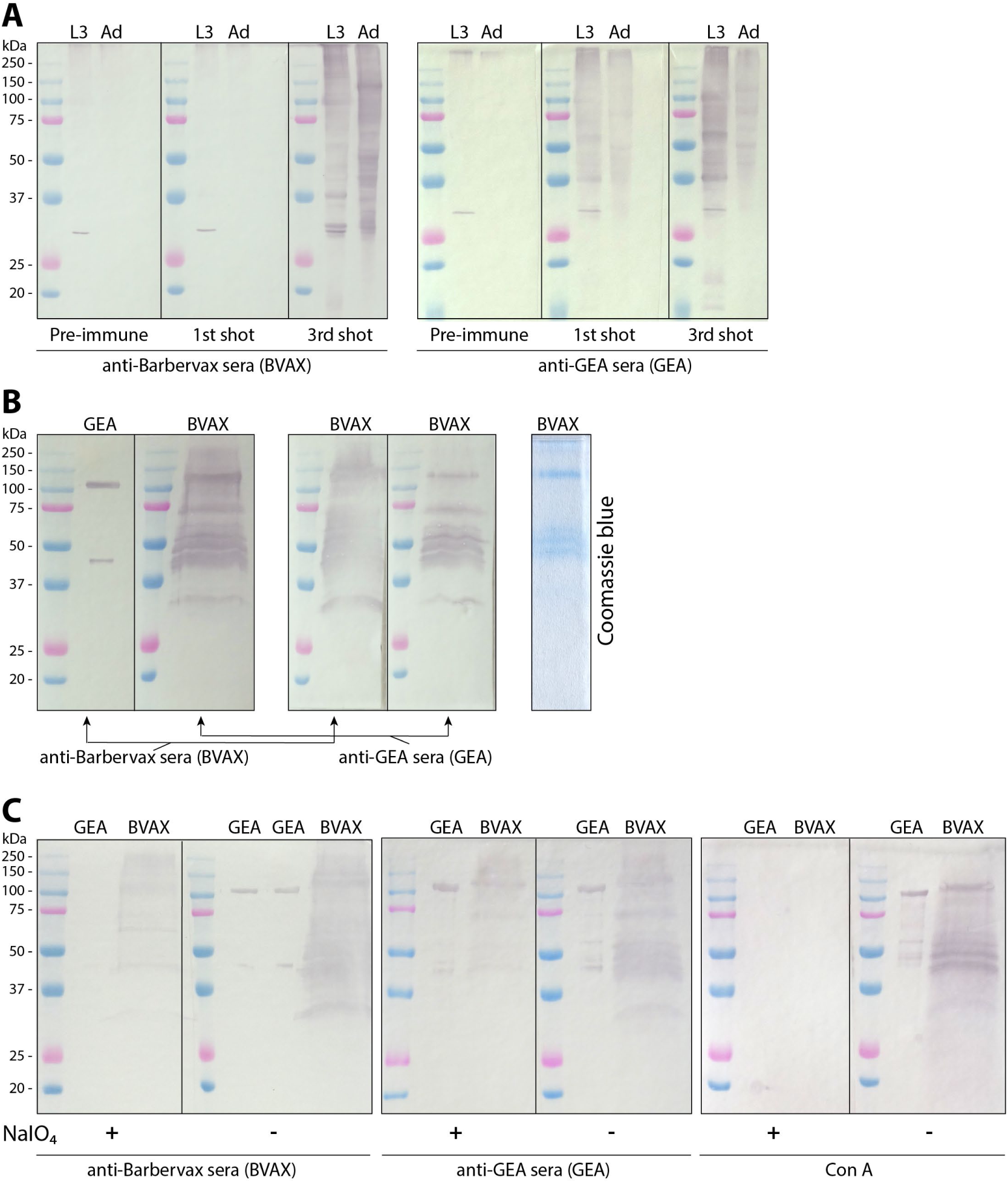
Recognition patterns of serum IgG antibodies to native and recombinant *H. contortus* antigens in vaccinated sheep. (A) vaccination with either Barbervax^®^ (BVAX) or GEA elevated IgG antibodies that bound to soluble proteins of L3 larvae (L3) and adult-stage (Ad) parasites. Parasite-specific IgG appeared earlier in GEA group than in BVAX group. (B) Cross-reactivities of IgG ware observed in both BVAX- and GEA-antisera; the comparison of Barbervax^®^-specific IgG (right panel) between the two groups suggested a similar binding pattern and intensity. (C) Deglycosylation of antigens using sodium metaperiodate (NaIO4) reduced Barbervax^®^-specific IgG targeting in both groups and completely abolished the recognition of GEA by BVAX IgG.

In addition to the appearance of IgGs, the binding spectrum of antibodies in animals vaccinated either with GEA or BVAX was compared. Antibodies in sera of animals vaccinated with GEA seemed to recognize many of the native antigens present in Barbervax^®^ (**Fig. 10B**). Indeed, the IgG recognition patterns towards Barbervax^®^ antigens displayed a comparable intensity and a similar molecular weight range. Similarly, sera from sheep vaccinated with Barbervax® showed comparable pattern recognition. Immunoblots were also performed on deglycosylated antigens, using Concanavalin A as a control to verify the completion of deglycosylation (**Fig. 10C**). IgG binding to the native antigens of Barbervax^®^ was decreased after metaperiodate treatment in both groups, while the recognition of GEA by BVAX IgG was completely abolished, strongly suggesting a glycan-biased targeting manner. In contrast to Barbervax^®^-induced IgG in the BVAX group, the intensity of the GEA group IgG towards deglycosylated GEA did not change.

#### Pathohistological examination

The pathohistological examination showed a cell infiltration of the lamina propria mucosae of the sheep in the BVAX and the GEA group. One ileum and one abomasum sample are shown as a representative for respective group (**Supplementary Fig. 1 and Fig. 2**). The ileum of the sheep from the BVAX group showed a higher cell density in the lamina propria mucosae in comparison to the other study groups.

## Discussion

The hematophagous parasite *H. contortus* is the most pathogenic trichostrongylid in small ruminants and South American camelids. Current control strategies primarily rely on the application of anthelmintic drugs, however the prevalence of anthelminthic resistance is emerging worldwide^8–10,13^. Barbervax^®^ has been the only commercialized vaccine against *H. contortus*, which high efficacy was reported in a number of controlled trials in sheep, goats and alpacas^18,29–31^. Despite this, Barbervax^®^ has two main drawbacks: First, it remains inaccessible in many countries due to biosecurity concerns, despite the fact that a safe, well-defined, and effective recombinant vaccine is urgently needed, second, there is a constant need to harvest antigen mixtures from worms retrieved from the stomachs of slaughtered sheep, as it is currently practiced, which makes it less attractive to ensure the adherence to 3R guidelines. Thus, a vaccine containing defined antigens that can be produced cost-effectively at large-scale in fermenters under controlled conditions, ensuring high hygiene and biosecurity standards, is urgently needed and would pave the way for the availability of vaccine doses year-round and worldwide. Suitable antigens have been indicated in studies using H11 and H-gal-GP antigens of the commercial vaccine Barbervax^®^. These studies further showed that these antigens are naturally glycosylated, carrying very distinct N-glycan structures compared to those found in mammals^32,33^. This made the production of recombinant forms of such antigens extremely challenging.

Previous studies using a combination of two recombinant H11 isoforms in Suffolk-cross lambs (antigen expressed in *C. elegans;* 3 injections with a saponin-based adjuvant; challenge with 5,000 L3) reported no significant difference in fecal egg shedding, worm burden or worm female: male ratio in comparison to the adjuvant control group^25^. While this study showed promising methodological results, it also became clear that correct glycosylation may play a far more important role than anticipated. Thus, in the present study, the safety and efficacy of two recombinant vaccines produced in Hi5 insect cells, one in a glycoengineered form (GEA), the other in non-glycoengineered form (NEA), were evaluated in a randomized controlled vaccine trial. To the authors’ best knowledge, the presented animal trial is the first successful attempt to vaccinate sheep with an antigen cocktail consisting of glycoengineered recombinant proteins produced in insect cells against a parasitic worm. Overall, both GEA and NEA did not show any appreciable negative side-effects in vaccinated animals. Both vaccines showed significant impact on the parasite, with egg shedding in the GEA group was reduced by 81.09% and worm burden decreased by 25.36%. A minimum efficacy of > 65% is required to have a positive epidemiological effect to control haemonchosis (*e.g.* reduced pasture contamination with subsequently lower worm burdens)^34^, and the presented GEA vaccine fulfilled this requirement in the present experimental trial with a wide margin. Despite a reduction of egg shedding by 32.72% in the NEA group compared to the unvaccinated group, no significant reduction in the worm burden was observed in this group. The comparison between GEA and NEA groups strongly indicated that glycoengineering is essential for an improved immune response against *H. contortus* since the GEA group continuously showed a more favorable outcome in parasitological and hematological parameters compared to the NEA group. Our data therefore confirm that nematode-type glycan modifications are crucial in the production of an anti-*Haemonchus* vaccine.

According to the manufacturer’s instructions for BVAX, the local tissue reactions in the form of swelling at the injection site can last for up to 17 days^35^. In the present study, the injection site was assessed on daily basis for seven days after each vaccination. There was no difference between groups apart form a transient increase of body temperature following the second vaccination which suggests that the insect cell-derived antigens are clinically safe.

*Haemonchus contortus* is hematophagous and can cause severe anemia in hosts^36^. As anemia in infected sheep is often accompanied by nutrient loss, either directly by the worms or indirectly due to the damaged intestinal epithelial layers, weight loss/regain after challenge was compared. Before challenge, there were no differences in daily weight gain between the groups, while after challenge the BVAX group showed the lowest body weight gain, which was somewhat unexpected, given that BVAX showed the highest reduction in worm burden/egg counts. Since many sheep are kept for meat production and body weight gain is a key performance indicator to measure the economic benefit of the farm, and differences in daily body weight gain were recorded after the challenge in the presenting study, it is suggested to include this parameter in future vaccine trials as an important criterion. Even though individual feed intake was not measured one can speculate that either daily dry matter intake was lower in the BVAX group (*e.g.* due to reduced appetite, abdominal pain) or that other mechanisms, such as an excessive immune reaction, caused the reduced daily weight gain. Indeed, vaccination against intestinal nematodes causes granulocyte infiltration in the mucosal layers. In previous works, during infection with trichostrongyles a potent chemoattractant activity for ovine bone marrow-derived eosinophils was noted *in vitro*^37^. This activity was identified as a chemoattractant protein in whole worm extracts of third and fourth larval and adult stages of *Teladorsagia circumcincta* as well as adult stages of *H. contortus*. The resulting local eosinophil-mediated mucosal damage, comparable to that seen in asthmatic lungs, may provide a permissive local microenvironment for the parasite for increased attachment and feeding^37,38^. Overall, eosinophils have been reported to play a significant role in gastrointestinal nematode infection in sheep^39^ and a negative association between blood eosinophils and fecal egg count (FEC) was observed in *H. contortus* infections^40^, and *in vitro* assays demonstrated that eosinophils may kill *H. contortus* larvae, thereby reducing parasite establishment^41^.

In the present study, pathohistological examination of the ileum of sheep in the BVAX group showed a higher density of cells in the lamina propria mucosae, compared to the other groups, which underscores this assumption. Additionally, the number of circulating eosinophils after challenge was higher in the BVAX group. It is therefore possible to speculate that the heightened immune response in the BVAX group, represented by higher concentrations of peripheral eosinophils and a higher cell density in the ileum, contributed to the reduced body weight gain. Furthermore, higher concentrations of eosinophils in the blood of sheep vaccinated with GEA compared to the blood of sheep vaccinated with NEA was observed. Following the 3^rd^ vaccination the eosinophils did not increase as drastically in all three experimental groups compared to the increase seen after the 1^st^ and the 2^nd^ injection. While pre-activation of eosinophils by vaccination may be an essential part of the induction of a protective immune response, subsequently limiting worm establishment, this assumption needs to be corroborated by analyzing the cytokine and chemokine profile in sheep sera generated from the different groups.

In addition to eosinophil concentration in peripheral blood, we also analyzed further blood parameters related to worm infection. Packed cell volume and hemoglobin levels can decrease severely after *H. contortus* infection since the sheep may lose up to 0.2 to 0.6 liters of blood per day^42^. In the present study, sheep vaccinated with GEA showed significantly higher total protein (5.95 g/dl versus 5.58 g/dl) and a numerically higher packed cell volume (26.77% versus 26.11%) compared to sheep vaccinated with NEA group. These results indicate a further benefit of the GEA vaccine that seems to significantly reduce blood losses through worm feeding. Even though the reduction in the worm burden in sheep vaccinated with GEA was only 25.36%, the PCV and Hb levels indicate that the remaining worms in GEA vaccinated animals were probably weakened. Whether this is part due to the impact of GEA on the integrity of the female worms, supported by the reduced number of female eggs per worm, remains to be followed up in detail in further trials. In this context, we also established whether the vaccine responses depended purely on secretory IgA, or whether a systemic adaptive immune response was involved. Our data suggested that IgG and IgE were the major isotypes that were elevated post vaccination, post infection as well as post-challenges. There was a link between parasite-specific antibodies and efficacy, with higher IgG/IgE level being associated with lower EpG values and worm burdens. The overall antibody titers specific for *Haemonchus* in the GEA group were lower than those in the BVAX group over the trial period, which could be the cause of a lower efficacy in this group.

The low titer of vaccine-specific antibodies observed in the BVAX group might be due to the low antigen coating efficiency despite optimization efforts. Independent of potential technical issues, ELISA data suggested that antibody in sera of sheep in the GEA group and NEA group increased quickly in comparison to the BVAX group, which indicated that the recombinant *H. contortus* antigens in the GEA and NEA preparation may be more efficient in inducing antigen-specific antibody responses compared to those being in the native worm antigens in Barbervax^®^.

Western blot analysis confirmed this observation and further revealed an unexpectedly broad antibody recognition pattern where numerous proteins from both L3 larvae and adult parasites were recognized by vaccine-induced antisera. Notably, IgG antibodies in sera of Barbervax^®^-vaccinated animals exhibited a strongly bias towards glycan moieties, which is in line with literature^25^. Similarly, anti-glycan IgG antibodies were detected in animals from the GEA group, alongside the ones specific to protein backbones. These findings implicate that the vaccine antigens, both the native and glycoengineered forms, can efficiently trigger antibody production in sheep that target a much broader range of glycoproteins than previously thought.

The novel glycoengineered vaccine GEA, comprising five glycoengineered recombinant antigens was tested for the first time *in vivo* in sheep and demonstrated that glycoengineering is essential to achieve a significant reduction in egg shedding and worm burden, and to improve hematological parameters. Although the overall efficacy of GEA at 100 µg/dose was lower than the commercial vaccine Barbervax^®^, the efficacy benchmark in this trial, animals in the GEA group exhibited greater body weight gain following challenge infection. Direct comparison, however, is complicated by the fact that Barbervax^®^ contains native H11 and additional components of the H-gal-GP complex, whereas GEA consists of a defined cocktail of glycoengineered recombinant H11 isoforms. However, it opens the way for future development of highly effective recombinant vaccines against this and other parasitic worms. To enhance the effectiveness of GEA, further studies to investigate the most effective recombinant antigen isoforms, the vaccine dose as well as immunization schedules are needed.

## Materials and methods

### Ethical considerations

The study was approved by the Ethics and Animal Welfare Committee of the University of Veterinary Medicine, Vienna in accordance with the University’s guidelines for Good Scientific Practice and authorized by the Austrian Federal Ministry of Education, Science and Research (vaccine trial: GZ:2023-0.734.950; *H. contortus* L3 production: GZ: 2022-0.599.404) in accordance with current legislation. The study was conducted according to the ARRIVE guidelines^43^.

### Sample size calculation and randomization

The sample size calculation was based on the following assumptions: vaccine efficacy is assessed by evaluating the number of excreted eggs (eggs per gram of feces; EpG) and the worm burden (number of worms in the abomasum). In the work by González-Sánchez et al. (2018) (publication Table 2), the standard deviation was approximately half of the mean of the total egg excretion, and the difference between the vaccinated and non-vaccinated groups was about 75%^44^. With a significance level of 5%, a power of 80%, and using a one-sided test procedure (vaccinated animals excrete fewer eggs per gram of feces), the calculated group size was 7 animals per group. The sheep were allocated to each study group randomly (stratified randomization) depending on their body weight and breed using the “rand()” function in Microsoft Excel.

### Sheep

The sheep were born between August and October 2023 (average weight of 26.7 kg; average age of 4 months) and raised on the same dairy sheep farm in the federal state of Lower Austria. The farm is run as a commercial organic dairy sheep farm with approximately 200 dairy sheep. The ewes were thoroughbred Lacaune and the three rams were thoroughbred Lacaune or Jura. Therefore, the lambs were Lacaune x Lacaune or Lacaune x Jura breed. The lambs were born and stayed with their mothers for 14 days before they were separated from the mother and kept in a separate barn. The lambs did not have any access to pasture, only to a concrete outdoor loafing area. The farmer separated 39 male lambs for the animal trial and housed them according to the national regulations. All lambs were vaccinated against clostridial disease on the farm within the regular herd health management procedure using Miloxan^®^ (Boehringer Ingelheim Animal Health France SCS, Lyon, France) with a dosage of 2 ml per sheep subcutaneously. They received the first immunization on October 25^th^, 2023, and the second one on November 28^th^, 2023.

### Deworming strategy before the trial

The first deworming was carried out on December 15^th^, 2023, where the sheep received 0.2 mg/kg body weight of ivermectin (Ivomec^®^ 10 mg/ml, Boehringer Ingelheim Animal Health) subcutaneously. At the farm of origin, the sheep were assessed, and the heaviest sheep was weighed and the dosage was chosen after that. Additionally, fecal samples were collected in September 2023 and examined for gastrointestinal nematodes, with negative results. At the day of arrival (January 2^nd^ and 3^rd^, 2024) the sheep were treated orally with 5 mg/kg BW of fenbendazole (Panacur^®^ 250 mg, Intervet GesmbH, Vienna), 7.5 mg/kg BW of netobimin (Hapadex^®^ 50 mg/kg, Intervet GesmbH, Vienna) and 20 mg/kg BW of toltrazuril (Baycox^®^ Multi 50mg/ml, Bayer Animal Health GmbH, Leverkusen, Germany).

### Vaccine efficacy trial in vivo

The randomized controlled vaccine trial was carried out at the Clinical Centre for Ruminant and Camelid Medicine, University of Veterinary Medicine, Vienna, Austria. The sheep were housed in a separate building and divided into five groups of seven animals (**Fig. 11**). All groups received the same amount of hay (not weighed) twice daily and water was provided *ad libitum*. In total, four groups were challenged with 5,000 in-house produced *H. contortus* L3 larvae, an isolate from Boehringer Ingelheim Vetmedica GmbH, Kathrinenhof Research Center, Rohrdorf, Bavaria, Germany. A negative control group (group NEG control; no vaccination and no challenge) and a positive control group (group POS control; no vaccination and challenge) were included. The 7 sheep in each experimental group were vaccinated three times subcutaneously either with 1 ml Barbervax^®^ (group BVAX; Wormvax Australia Pty Ltd., Albany, WA, Australia; imported from UK via Merlin Vet, Kelso, UK), or with 1 ml of the glycoengineered recombinant antigen cocktail (group GEA,^28^) or with 1 ml of the non-glycoengineered recombinant antigen cocktail (group NEA). Recombinant vaccines (GEA and NEA) were formulated by mixing 100 µg of insect cell (Hi5)-derived recombinant products (H11, H11-1, H11-2, H11-4 and GA1; 20 µg/antigen/dose) with 1 mg of Quil-A^®^ adjuvant (InvivoGen, San Diego, CA, USA) in PBS solution. Sheep were challenged one week after the third vaccination using a 2 ml disposable plastic pipette (dosage 2 ml). Multiple samples, including fecal, blood and saliva samples, were collected from animals during the study period (**Fig. 11**)

**Figure 11:**
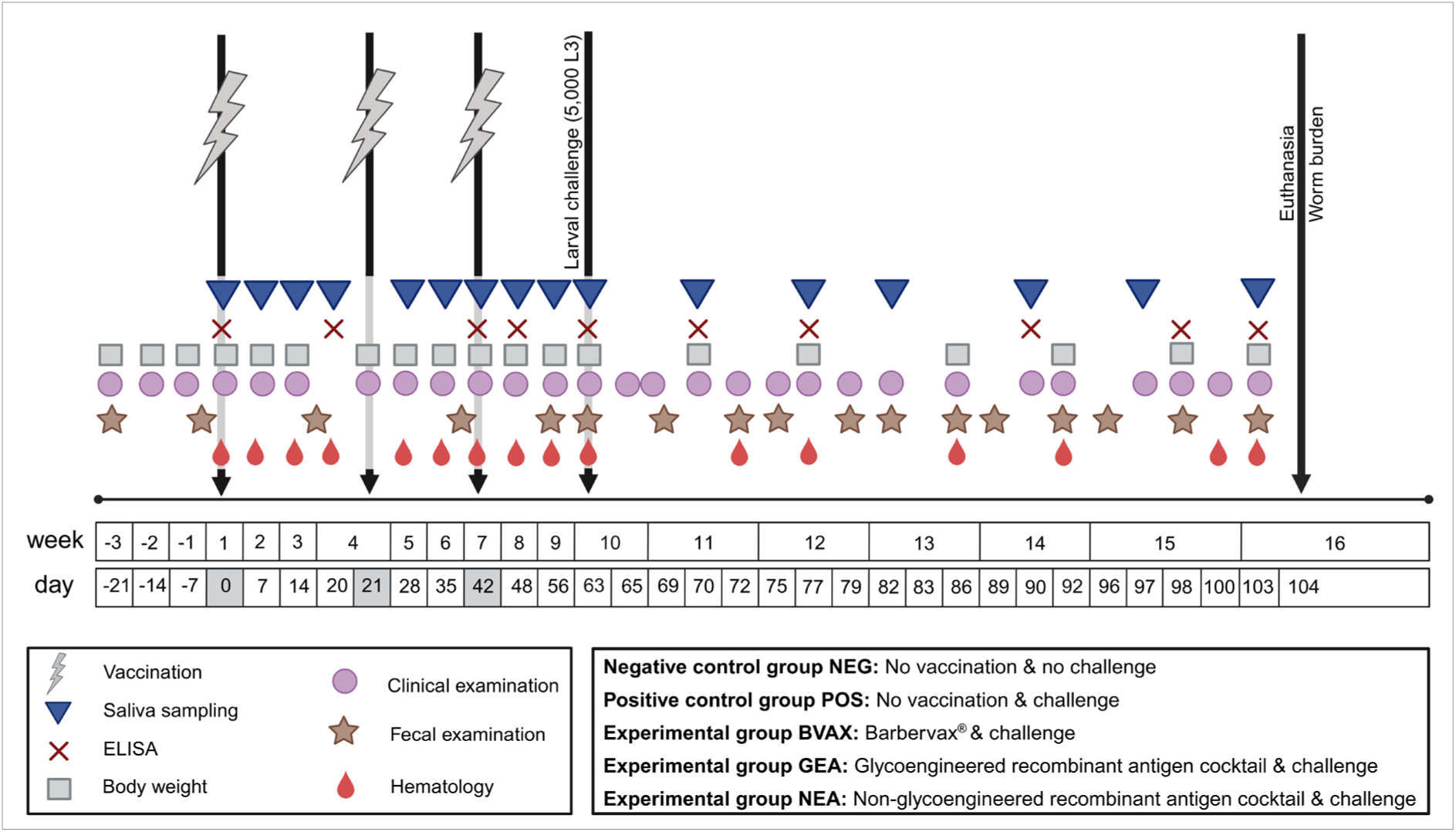
Overview on the specimen material collected during the vaccine trial from Jan to May 2024. The sheep were kept in five groups with seven sheep in each group. The negative control group (NEG) received no vaccination and no challenge. The positive control (POS) and the experimental groups were challenged with 5,000 *H. contortus* L3 (in week 10). The experimental groups were vaccinated three times subcutaneously with Barbervax^®^ (BVAX), the novel glycoengineered vaccine (GEA) or the non-glycoengineered vaccine (NEA). Examinations, sampling dates and ELISA test dates are shown [*Created in BioRender. Sajovitz-Grohmann, F. (2025)* https://BioRender.com/pkpssk5].

The production of the glycoengineered recombinant antigen cocktail is patented under the intellectual property IP (European Patent Application No. EP24216456.4 entitled “Expression system for glycoengineered antigens” filed November 29, 2024). Detailed description of the production and *in vitro* characterization of the antigen cocktail is described by Adduci et al., 2025^28^.

### Clinical assessment and differential blood counts

The clinical assessment (e.g. body temperature, FAMACHA^©^) was carried out using a standardized protocol for physical examination and blood samples (EDTA, serum, Tempus™ Blood RNA Tubes [Thermo Fisher Scientific, Waltham, MA, USA]) were drawn from the external jugular vein at defined time points throughout the study period (**Fig. 11**)^45–47^. The sheep were weighed using Gallagher W0 digital scales (Gallagher Europe, Groningen, The Netherlands). Hematological parameters of whole blood sample, including leucocytes, erythrocytes, hemoglobin, and erythrocyte indices, were analyzed by flow cytometry (Advia^®^ 2120i, Siemens, Erlangen, Germany). Total protein and albumin were measured by photometry (Cobas^®^ pure C303, Roche Diagnostics International Ltd, Rotkreuz, Switzerland). During the whole study period, the vaccination was carried out after the clinical assessment and blood sample collection.

### Assessment after vaccination and local injection site reaction

The 21 sheep from the BVAX, GEA and NEA groups were assessed eight times after each vaccination. The local injection site was assessed on the day of vaccination (morning: vaccination; afternoon: first assessment) until day 7 after vaccination once a day following the the injection site and the surrounding area were assessed: swelling (none; slight; severe), pain during palpation (yes/no), necrosis (none; slight; severe), tissue induration (none; firm; hard) and lymph node enlargement (none; slight; severe).

#### Preparation of soluble H. contortus proteins

Soluble proteins from adult *H. contortus* were extracted using a two-step homogenization process. Briefly, adult worms were resuspended in Tris-buffered saline (100 mM Tris-HCl, 100 mM NaCl, pH 7.4) supplemented with 0.02% sodium azide, 1% octylthioglucoside, and a 1% protease inhibitor cocktail (Sigma-Aldrich, St. Louis, MO, USA). The worms were first homogenized on ice for 2 min with intermittent pauses using a Turrax^®^ dispersion tool (T10 basic, equipped with an S10N-10G-ST probe; IKA, Staufen, Germany). The homogenate was then transferred to an ice-cold Dounce homogenizer for manual homogenization. Resulted homogenate was centrifuged at 12,000 × *g* for 30 min at 4 °C. The supernatant was filtered through 0.22 µm sterile syringe filters and collected into sterile Falcon tubes. For larval (L3) parasites, due to the limited material, homogenization was performed in 1.5 ml tubes using disposable polypropylene pestles (Thermo Fisher Scientific). Protein concentration was measured using a Pierce™ Bradford Plus protein assay kit (Thermo Fisher Scientific).

#### Enzyme-linked immunosorbent assay (ELISA)

Native antigens of adult *H. contortus* and recombinant antigen cocktails (GEA and NEA) were diluted to a concentration of 2 μg/ml in coating buffer (200 mM carbonate/bicarbonate, pH 9.6). 96-well Nunc-Immuno™ MicroWell™ ELISA plates (Sigma-Aldrich) were coated overnight at 4 °C with 50 μl of antigens (100 ng/well). Post blocking with 2% BSA, sheep sera (dilution 1:100) were added to the wells and incubated at 37 °C for 1 hour. Serum antibodies were probed with anti-sheep IgG (Sigma-Aldrich, dilution 1:10,000), IgE (Bio-Rad Laboratories Ges.m.b.H., Hercules, CA, USA; 1E7, dilution 1:10,000), IgM (Sigma-Aldrich; rabbit polyclonal, dilution 1:10,000) and IgA (Bio-Rad Laboratories Ges.m.b.H; rabbit polyclonal, dilution 1:1,000). Horseradish peroxidase(HRP)-conjugated secondary antibodies, anti-mouse IgG (Sigma-Aldrich, dilution 1:5,000) and anti-rabbit IgG (Sigma-Aldrich dilution 1:10,000), were used to react with tetramethylbenzidine-hydrogen peroxide (Thermo Fisher Scientific). Post addition of a reaction stopping buffer (Carl Roth GmbH + Co. KG, Karlsruhe, Germany; 0.5 M sulfuric acid), optical density at 450 nm (OD450) was measured using a microplate reader (FilterMax F5; Molecular Devices, San Jose, CA, USA). Total serum IgG and IgM were examined by performing direct ELISA using an anti-sheep antibody (stated above) to probe sheep sera (1:100 diluted) coated into ELISA plates.

#### Western blotting and metaperiodate treatment

Worm-derived and recombinant antigens were dissolved in SDS-loading dye, denatured at 95 °C and loaded on 12% SDS-PAGE gels. Precision Plus Protein™ Dual Color Standards (Bio-Rad Laboratories GesmbH) was used as protein ladder. Post electrophoresis at 200 V, proteins were transferred onto nitrocellulose membranes using a TransBlot^®^ Turbo™ Transfer System (Bio-Rad Laboratories GesmbH). After blocking in 0.5% BSA, nitrocellulose membranes were incubated with sheep antisera, combined from the 7 sheep of the same group (dilution 1:200), then with a mouse anti-goat/sheep IgG (Sigma-Aldrich; dilution 1:10,000), and finally with an alkaline phosphatase (AP)-conjugated anti-mouse IgG (Sigma-Aldrich; Fc-specific; dilution 1:10,000). SIGMAFAST™ BCIP^®^/NBT (Sigma Aldrich) was used as the substrate for color development.

Protein deglycosylation was performed on-membrane using a protocol adapted from Roberts et al.^25^. Briefly, immunoblots were treated overnight at 4 °C with 10 mM sodium meta-periodate dissolved in 0.05 M sodium acetate, pH 4.5, 0.05 M sodium chloride buffer, prior to subsequent blocking and incubation steps. To monitor the deglycosylation efficiency, a treated blot was incubated with biotinylated Concanavalin A (Vector Laboratories, Newark, CA, USA; Con A, dilution 1:250) in PBS buffer supplemented with 1 mM calcium chloride followed by a goat anti-biotin secondary antibody (Sigma-Aldrich; dilution 1:10,000).

#### Efficacy assessment Mini-FLOTAC method

Feces were sampled and analyzed individually by Mini-FLOTAC with a detection limit of five eggs per gram (EpG)^49,50^. Samples were weighed (5 g feces) and mixed with 45 ml of a saturated saline solution (density 1.18 g/ml) and sieved through a mesh (average mesh size: 1.3 mm) in a beaker afterwards. An equal distribution of the eggs was ensured by using a magnetic stirrer (IKA-COMBIMAG REO, Janke & Kunkel GmbH u. Co. KG, Staufen, Germany). Samples were quantitatively evaluated under the light microscope (Eclipse Ci-S, 100 x magnification, Nikon, Vienna, Austria). The EpG was calculated by multiplying the number of eggs counted in both chambers by 5 (multiplication factor for livestock feces)^50^.

#### Worm burden and worm sexing

The worm burden was evaluated after euthanizing the sheep. First, the abdominal cavity was opened, and the duodenum was ligated. The omasum was cut away to keep the content within the abomasum. Subsequently, the abomasum was removed from the gastro-intestinal bundle and put into a bucket. The greater curvature was opened, and the content was transferred into a sieve with a mesh width of 150 µm and washed thoroughly. Subsequently, the entire abomasal content was transferred into a bucket for storage and further assessment. The abomasal mucosa as well as the entire abomasal content were inspected for adult worms under good light conditions. All worms were collected, separated by gender and counted within three days after sacrifice^51,52^.

#### Patho-histological examination

Abomasal and ileum tissues were collected for pathohistological examination *postmortem*. Briefly, abomasal tissue (location fundus region) and ileum tissue, in size of one square-centimeter, were taken immediately after opening the abdominal cavity using a scissor and forceps and fixed in 10% formalin (BiopSafe^®^, Vedbœk, Denmark) in a 20 ml container. Samples were collected from three sheep from each group. Subsequently, the samples were dehydrated in graded series of ethanol and embedded in paraffin wax. Thereafter, approximately 3 μm thick sections were stained with HE for further assessment.

#### Statistics

Descriptive and explorative statistical analysis was performed using Microsoft Excel 2010 (Microsoft, Washington, USA) and IMB SPSS^®^ Statistics Version 29 (IBM, New York, USA). Descriptive statistics were carried out using mean, median, 25^th^ and 75^th^ percentile, minimum and maximum. Normal distribution was tested using the Kolmogorov-Smirnov test. The daily weight gain in gram was calculated before (63 days) and after (40 days) challenge by using the body weight at the begin of the trial (2^nd^/3^rd^ January), before challenge (27^th^ March) and at the end of the trial (6^th^ May). For the packed cell volume, thresholds were calculated using the 25^th^, 50^th^ and 75^th^ percentile. The cumulative egg excretion was calculated by summarizing the egg excretions over time. The EpG values per worm was calculated by dividing the cumulative EpG values by the number of female worms (no normal distribution). The female/male ratio was calculated by dividing female by male worms. The EpG values and worm counts were log transformed using the lg10 function in SPSS. All measures (EpG values, worm counts, blood values) were analyzed using a linear mixed model followed by a post hoc test (Sidak test) between the experimental groups. Group identification and sampling date were included as fixed values. Sampling date was included as repeated measure and sheep identification as subject. All missing values were excluded in the analysis. The Akaike’s information criterion (AIC) revealed 215.24 for the egg excretion. Comparison between groups without repeated measures was carried out using the Kruskal Wallis test (> 2 samples) including the Bonferroni correction (e.g. worm burden, cumulative EpG), the Mann-Whitney U test (2 samples) or the Fishers exact test (< 5 cases). For comparison between groups with a normal distribution a one-way ANOVA was carried out. For all tests the significance level was set at p ≤ 0.05.

## Data availability

The datasets generated during and/or analyzed during the current study are available from the corresponding author on reasonable request.

## Acknowledgements

We thank Drs Dietmar Hamel and Steffen Rehbein for sharing their expertise at multiple occasions during the whole project period. Special thanks go to the students who supported the whole team during the trial, especially Gudrun Gerstner, Judith Hauth, Vinzent Leonfellner, Roman Rupprechter, Christina Schroll, Vanessa Stadlmann, Regina Walch and Glenn Wang. Additionally, we thank Roman Peschke, Sonja Rohrer, Bärbel Ruttkowski, Maria Unterköfler, and Hugo Weidinger at the Institute of Parasitology who supported us in the laboratory and the Clinical Pathology Platform who performed the hematological examinations. Special thanks go to René Brunthaler who supported us in preparing the pathology images.

This project has been funded by the Program "Top Vet Science" financed by Vetmeduni Vienna.

## Author contributions statement

S.Y., K.L., A.J., T.W. and D.W. conceived the project. S.Y., K.L., A.J., T.W., F.SG., I.A., B.H. and D.W. contributed to the design of the methodology. S.Y., K.L., F.SG., I.A., D.W. and A.T. carried out the formal analysis. S.Y., K.L., F.SG., I.A., J.E., J.Z., B.P., L.W., S.W. and B.H. performed the experiments and collected the data. A.J., T.W., B.H., S.Y., K.L., F.SG., I.A., S.W., and L.W. provided the resources in terms of labor resources, materials, reagents and analysis tools. F.SG. and K.L. prepared the initial draft of the manuscript. F.SG., K.L. and I.A. visualized the data. K.L., S.Y., T.W., A.J., B.H. and D.W. acted as supervisors. S.Y., K.L., F.SG and I.A. managed the project administration. S.Y., K.L., T.W. and A.J. acquired the funding. F.SG., K.L., S.Y., A.J., D.W., T.W., I.A., B.H., L.W., B.P., S.W., J.Z., J.E. and A.T. reviewed and edited the manuscript and accepted the final version of the manuscript.

