## Supplementary File for "Safety and Efficacy of a Novel Glycoengineered Recombinant Vaccine Candidate against *Haemonchus contortus* in Sheep"

**Supplementary Table 1**: Injection site assessment after the first vaccination. The assessment was carried out daily, starting on the day of the vaccination (induration/swelling/lymph node enlargement) until day 7 after the vaccination. The table shows the grade of severity of induration and swelling of the injection site and lymph node enlargement (not detected, slight, severe) in the sheep from the BVAX, GEA and NEA group.

| | | **Induration** | | | **Total** |
|---|---|---|---|---|---|
| | | **no** | **slight** | **severe** | |
| **Group ID** | BVAX | 10 | 21 | 25 | 56 |
| | GEA | 11 | 19 | 26 | 56 |
| | NEA | 2 | 23 | 31 | 56 |
| **Total** | | 23 | 63 | 82 | 168 |
| | | **Lymph node enlargement** | | | |
| | | **no** | **slight** | **severe** | |
| **Group ID** | BVAX | 10 | 33 | 13 | 56 |
| | GEA | 1 | 21 | 34 | 56 |
| | NEA | 2 | 22 | 32 | 56 |
| **Total** | | 13 | 76 | 79 | 168 |
| | | **Swelling** | | | |
| | | **no** | **slight** | **severe** | |
| **Group ID** | BVAX | 6 | 35 | 15 | 56 |
| | GEA | 1 | 31 | 24 | 56 |
| | NEA | 0 | 37 | 19 | 56 |
| **Total** | | 7 | 103 | 58 | 168 |

**Supplementary Table 2**: Injection site assessment after the second vaccination. The assessment was carried out daily, starting on the day of the vaccination (induration/swelling/lymph node enlargement) until day 7 after the vaccination. The table shows the grade of severity of induration and swelling of the injection site and lymph node enlargement (not detected, slight, severe) in the sheep from the BVAX, GEA and NEA group.

| | | Induration | | | Total |
|---|---|---|---|---|---|
| | | **no** | **slight** | **severe** | |
| **Group ID** | BVAX | 22 | 22 | 12 | 56 |
| | GEA | 4 | 8 | 44 | 56 |
| | NEA | 19 | 16 | 21 | 56 |
| **Total** | | 45 | 46 | 77 | 168 |
| | | **Lymph node enlargement** | | | |
| | | **no** | **slight** | **severe** | |
| **Group ID** | BVAX | 22 | 22 | 12 | 56 |
| | GEA | 3 | 19 | 34 | 56 |
| | NEA | 13 | 21 | 22 | 56 |
| **Total** | | 38 | 62 | 68 | 168 |
| | | **Swelling** | | | |
| | | **no** | **slight** | **severe** | |
| **Group ID** | BVAX | 16 | 29 | 11 | 56 |
| | GEA | 7 | 21 | 28 | 56 |
| | NEA | 13 | 32 | 11 | 56 |
| **Total** | | 36 | 82 | 50 | 168 |

**Supplementary Table 3**: Injection site assessment after the third vaccination. The assessment was carried out daily, starting on the day of the vaccination (induration/swelling/lymph node enlargement) until day 7 after the vaccination. The table shows the grade of severity of induration and swelling of the injection site and lymph node enlargement (not detected, slight, severe) in the sheep from the BVAX, GEA and NEA group.

| | | Induration | | | Total |
|---|---|---|---|---|---|
| | | **no** | **slight** | **severe** | |
| **Group ID** | BVAX | 19 | 13 | 24 | 56 |
| | GEA | 4 | 8 | 44 | 56 |
| | NEA | 11 | 21 | 24 | 56 |
| **Total** | | 34 | 42 | 92 | 168 |
| | | **Lymph node enlargement** | | | |
| | | **no** | **slight** | **severe** | |
| **Group ID** | BVAX | 11 | 15 | 30 | 56 |
| | GEA | 2 | 13 | 41 | 56 |
| | NEA | 5 | 23 | 28 | 56 |
| **Total** | | 18 | 51 | 99 | 168 |
| | | **Swelling** | | | |
| | | **no** | **slight** | **severe** | |
| **Group ID** | BVAX | 17 | 34 | 5 | 56 |
| | GEA | 2 | 37 | 17 | 56 |
| | NEA | 13 | 41 | 2 | 56 |
| **Total** | | 32 | 112 | 24 | 168 |

**Supplementary Table 4**: Injection pain assessment of the infection site after the first, second and third vaccination was evaluated. The assessment was carried out daily, starting on the day of the vaccination until day 7 after the vaccination. The table shows the results of the evaluation (painful yes/ painful no) in the sheep from the BVAX, GEA and NEA group.

| | | Painful | | Total |
|---|---|---|---|---|
| | | no | yes | |
| | **Vaccination 1** | | | |
| | BVAX | 22 | 34 | 56 |
| **Group ID** | GEA | 16 | 40 | 56 |
| | NEA | 6 | 50 | 56 |
| **Total** | | 44 | 124 | 168 |
| | **Vaccination 2** | | | |
| | BVAX | 44 | 12 | 56 |
| **Group ID** | GEA | 25 | 31 | 56 |
| | NEA | 31 | 25 | 56 |
| **Total** | | 99 | 68 | 167 |
| | **Vaccination 3** | | | |
| | BVAX | 31 | 25 | 56 |
| **Group ID** | GEA | 21 | 35 | 56 |
| | NEA | 30 | 26 | 56 |
| **Total** | | 82 | 86 | 168 |

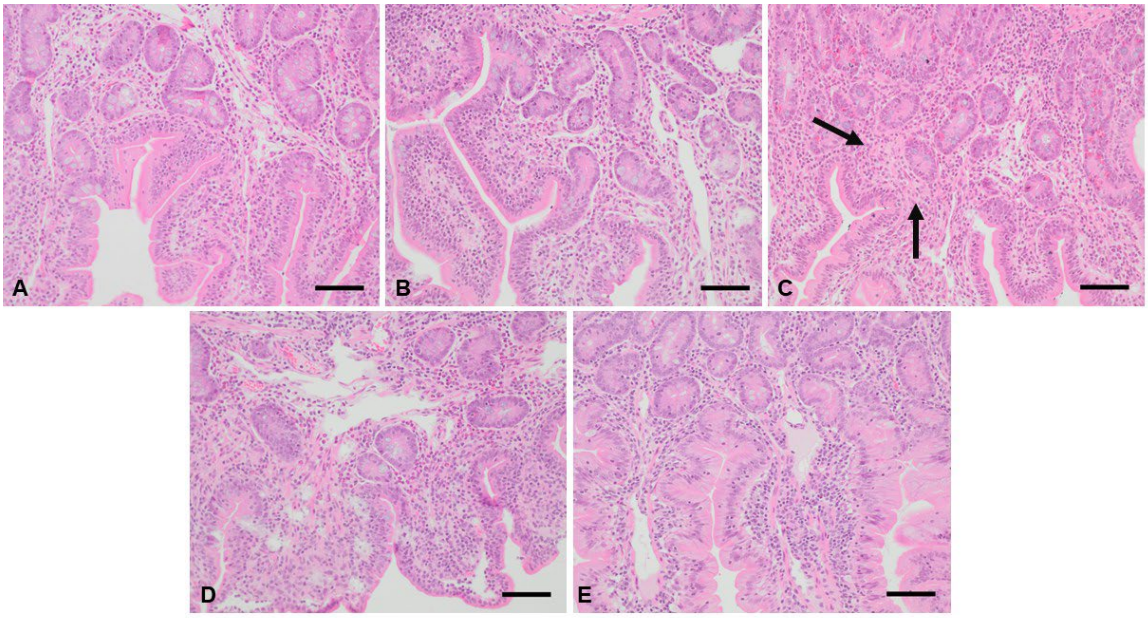


**Supplementary Figure 1:** Ileum following hematoxylin/eosin (HE) staining (200-fold magnification; bar: 80 µm). A) sheep VT3 from the NEG group, B) sheep VT8 from the POS group, C) VT16 from the BVAX group, D) sheep VT24 from the GEA group and E) sheep VT31 from the NEA group. One ileum sample for respective group is shown as a representative.

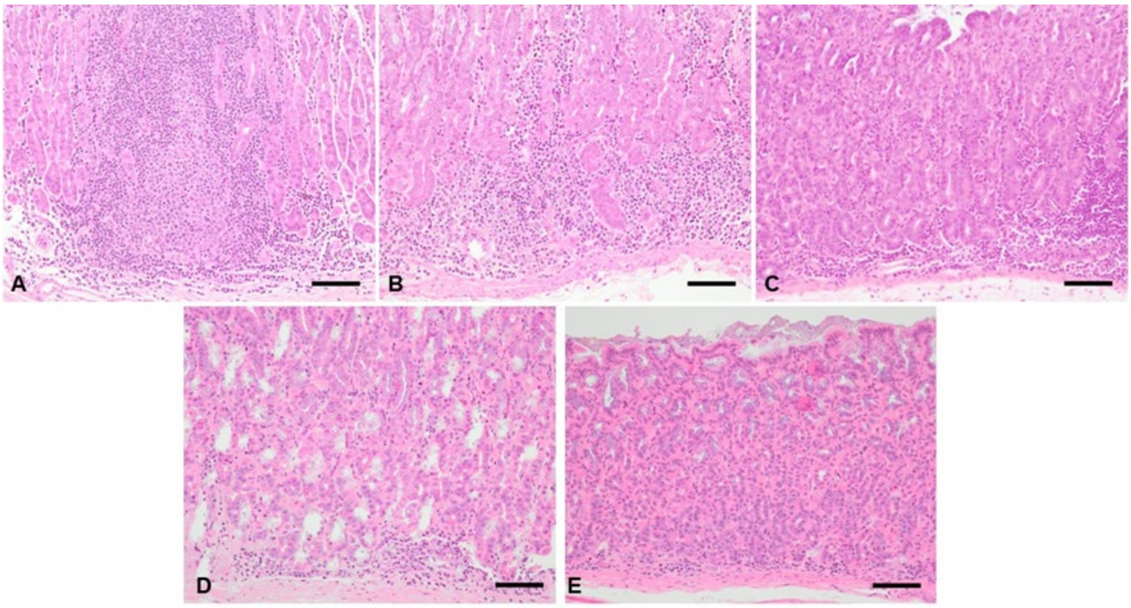


**Supplementary Figure 2:** Abomasal tissue samples from the fundus region following Haematoxylin/Eosin staining (200-fold magnification; bar = 200 µm). A) sheep VT3 from the negative control group, B) sheep VT8 from the positive control group, C) sheep VT 16 from the BVAX group, D) sheep VT24 from the GEA group and E) sheep VT31 from the NEA group.

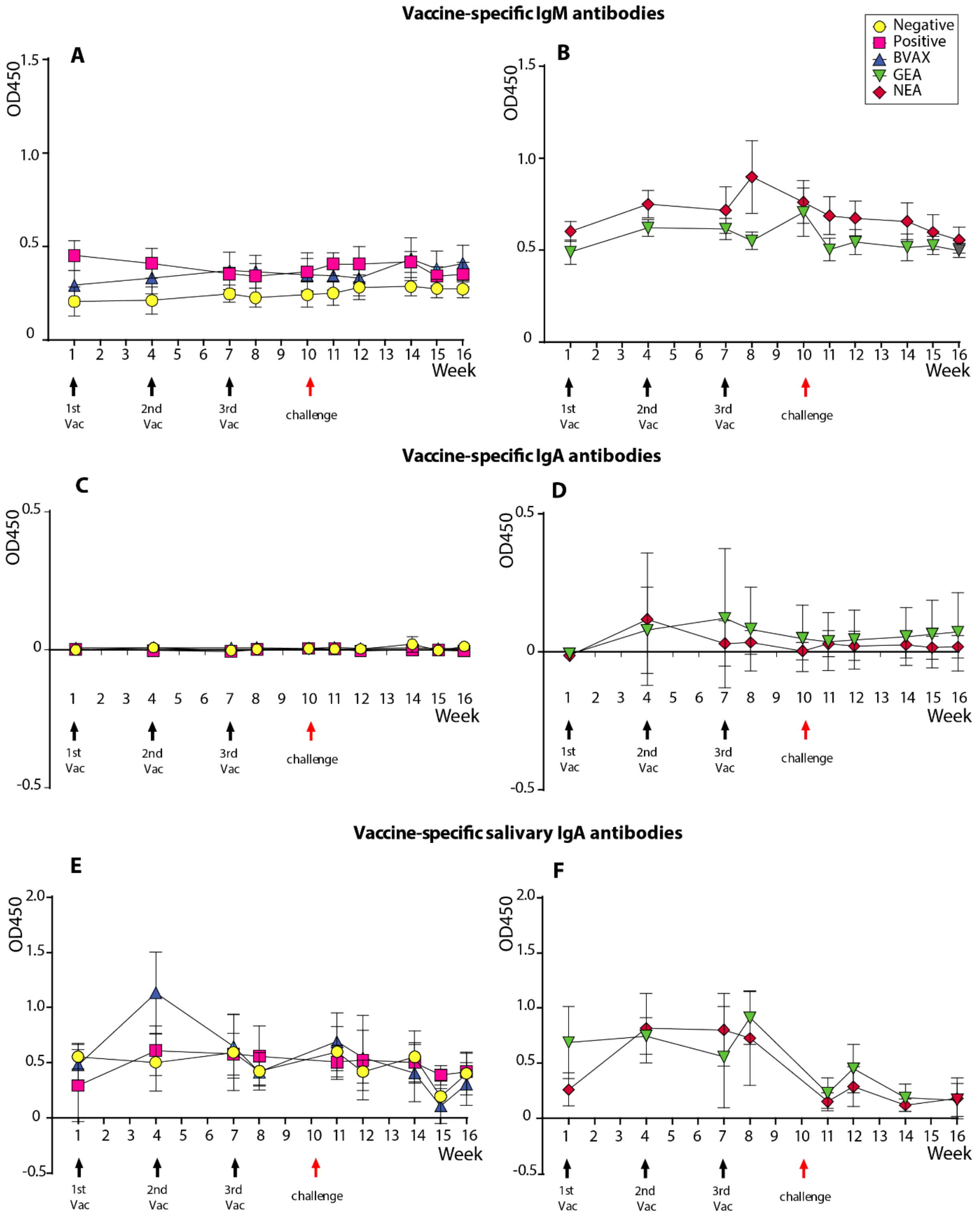


**Supplementary Figure 3. ELISA analysis of serum IgM, IgA and salivary IgA antibodies in sheep.** Panels (**A-D**) depict antigen-specific IgM and IgA responses, respectively. Panels (**E, F**) illustrate salivary IgA responses. Data represent individual sheep with the average shown. Vaccine-specific antibodies were measured using plates coated with equal amount of corresponding vaccine antigens, either with Barbervax®, or GEA or NEA. Control groups (negative and positive) were assayed using Barbervax®-coated plates.
